# Deep parallel characterization of AAV tropism and AAV-mediated transcriptional changes via single-cell RNA sequencing

**DOI:** 10.1101/2021.06.25.449955

**Authors:** David Brown, Michael Altermatt, Tatyana Dobreva, Sisi Chen, Alexander Wang, Matt Thomson, Viviana Gradinaru

**Affiliations:** Division of Biology and Biological Engineering, California Institute of Technology, Pasadena, CA, USA; Andrew and Peggy Cherng Department of Medical Engineering, California Institute of Technology, Pasadena, CA, USA; Division of Engineering and Applied Science, California Institute of Technology, Pasadena, CA, USA

## Abstract

Engineered variants of recombinant adeno-associated viruses (rAAVs) are being developed rapidly to meet the need for gene-therapy delivery vehicles with particular cell-type and tissue tropisms. While high-throughput AAV engineering and selection methods have generated numerous variants, subsequent tropism and response characterization have remained low throughput and lack resolution across the many relevant cell and tissue types. To fully leverage the output of these large screening paradigms across multiple targets, we have developed an experimental and computational single-cell RNA sequencing (scRNA-seq) pipeline for in vivo characterization of barcoded rAAV pools at unprecedented resolution. Using our platform, we have corroborated previously reported viral tropisms and discovered unidentified AAV capsid targeting biases. As expected, we observed that the tropism profile of AAV.CAP-B10 in mice was shifted toward neurons and away from astrocytes when compared with AAV-PHP.eB. Our transcriptomic analysis revealed that this neuronal bias is mainly due to increased targeting efficiency for glutamatergic neurons, which we confirmed by RNA fluorescence in situ hybridization. We further uncovered cell subtype tropisms of AAV variants in vascular and glial cells, such as low transduction of pericytes and Myoc+ astrocytes. Additionally, we have observed cell-type-specific responses to systemic AAV-PHP.eB administration, such as upregulation of genes involved in p53 signaling in endothelial cells three days post-injection, which return to control levels by day twenty-five. Such ability to parallelize the characterization of AAV tropism and simultaneously measure the transcriptional response of transduction will facilitate the advancement of safe and precise gene delivery vehicles.

## Introduction

Recombinant AAVs (rAAVs) have become the preferred gene delivery vehicles for many clinical and research applications (Bedbrook et al., 2018; Samulski and Muzyczka, 2014) owing to their broad viral tropism, ability to transduce dividing and non-dividing cells, low immunogenicity, and stable persistence as episomal DNA ensuring long-term transgene expression (Daya and Berns, 2008; Deverman et al., 2018; Gaj et al., 2016; Hirsch and Samulski, 2014; Naso et al., 2017; Wu et al., 2006). However, current systemic gene therapies using AAVs have a relatively low therapeutic index (Mével et al., 2020). High doses are necessary to achieve sufficient transgene expression in target cell populations, which can lead to severe adverse effects from off-target expression (Hinderer et al., 2018; Srivastava, 2020; Wilson and Flotte, 2020). Increased target specificity of rAAVs would reduce both the necessary viral dose and off-target effects: thus, there is an urgent need for AAV gene delivery vectors that are optimized for cell-type-specific delivery (Paulk, 2020). Lower viral doses would also alleviate demands on vector manufacturing and minimize the chances of undesirable immunological responses (Calcedo et al., 2018; Gao et al., 2009; Mingozzi and High, 2013). Capsid-specific T-cell activation was reported to be dose-dependent in vitro (Finn et al., 2010; Pien et al., 2009) and in humans (Mingozzi et al., 2009; Nathwani et al., 2011). Shaping the tropism of existing AAVs to the needs of a specific disease has the potential to reduce activation of the immune system by detargeting cell types, such as dendritic cells, that have an increased ability to activate T-cells (Herzog et al., 2019; Rogers et al., 2017; Rossi et al., 2019; Somanathan et al., 2010; Vandenberghe et al., 2006; Zhu et al., 2009).

Several studies have demonstrated that the transduction efficiency and specificity of natural AAVs can be improved by engineering their capsids using rational design (Bartlett et al., 1999; Davidsson et al., 2019; Davis et al., 2015; Lee et al., 2018; Sen, 2014) or directed evolution (Chan et al., 2017; Dalkara et al., 2013; Deverman et al., 2016; Excoffon et al., 2009; Grimm et al., 2008; Körbelin et al., 2016a; Kotterman and Schaffer, 2014; Maheshri et al., 2006; Müller et al., 2003; Ogden et al., 2019; Ojala et al., 2018; Pekrun et al., 2019; Pulicherla et al., 2011; Ravindra Kumar et al., 2020; Tervo et al., 2016; Ying et al., 2010). These engineering methods yield diverse candidates that require thorough, preferably high-throughput, in vivo vector characterization to identify optimal candidates for a particular clinical or research application. Toward this end, conventional immunohistochemistry (IHC) and various in situ hybridization (ISH) techniques are commonly employed to profile viral tropism by labeling proteins expressed by the viral transgene or viral nucleic acids, respectively (Arruda et al., 2001; Chan et al., 2017; Deleage et al., 2016, 2018; Deverman et al., 2016; Grabinski et al., 2015; Hinderer et al., 2018; Hunter et al., 2019; Miao et al., 2000; Polinski et al., 2015, 2016; Puray-Chavez et al., 2017; Ravindra Kumar et al., 2020; Wang et al., 2020; Zhang et al., 2016; Zhao et al., 2020).

Although these histological approaches preserve spatial information, current technical challenges limit their application to profiling the viral tropism of just one or two AAV variants across a few gene markers, thus falling short of efficiently characterizing multiple AAVs across many complex cell types characteristic of tissues in the central nervous system (CNS). The reliance on known marker genes also prevents the unbiased discovery of tropisms since such marker genes need to be chosen *a priori*. Choosing marker genes is particularly challenging for supporting cell types, such as pericytes in the CNS microvasculature and oligodendrocytes, which often have less established cell type identification strategies (Liu et al., 2020; Marques et al., 2016). The advent of single-cell RNA sequencing (scRNA-seq) has enabled comprehensive transcriptomic analysis of entire cell-type hierarchies, and brought new appreciation to the role of cell subtypes in disease (Berto et al., 2020; Gokce et al., 2016; Tasic et al., 2016, 2018; Zeisel et al., 2018). However, experimental and computational challenges, such as the sparsity of RNA capture and detection, strong batch effects between samples, and the presence of ambient RNA in droplets, reduce the statistical confidence of claims about individual gene expression (Lähnemann et al., 2020; Yang et al., 2020; Zheng et al., 2017). Computational methods have been developed to address some of these challenges, such as identifying contaminating RNA (Yang et al., 2020), accounting for or removing batch effects (Korsunsky et al., 2019; Lin et al., 2019; Lopez et al., 2018), and distinguishing intact cells from empty droplets (Lun et al., 2019; Macosko et al., 2015; Zheng et al., 2017). However, strategies for simultaneously processing transcripts from multiple delivery vehicles and overcoming the computational challenges of confidently detecting individual transcripts have not yet been developed for probing the tropism of AAVs in complex, heterogeneous cell populations.

Collecting the entire transcriptome of injected and non-injected animals offers an opportunity to study the effects of AAV transduction on the host cell transcriptome. A similar investigation has been conducted with G-deleted rabies virus (Huang and Sabatini, 2020). This study demonstrated that virus infection led to the downregulation of genes involved in metabolic processes and neurotransmission in host cells, whereas genes related to cytokine signaling and the adaptive immune system were upregulated. At present, no such detailed examination of transcriptome changes upon systemic AAV injection has been conducted. High-throughput single-cell transcriptomic analysis could provide further insight into the ramifications of AAV capsid and transgene modifications with regard to innate (Duan, 2018; Hösel et al., 2012; Martino et al., 2011; Shao et al., 2018; Zaiss et al., 2008) and adaptive immune recognition (George et al., 2017; Manno et al., 2006; Mingozzi et al., 2007; Nathwani et al., 2011, 2014). Innate and adaptive immune responses to AAV gene delivery vectors and transgene products constitute substantial hurdles to their clinical development (Colella et al., 2018; Shirley et al., 2020). The study of brain immune response to viral gene therapy has been limited to antibody staining and observation of brain tissue slices post direct injection. In particular, prior studies have shown that intracerebral injection of rAAV vectors in rat brains does not induce leukocytic infiltration or gliosis (Chamberlin et al., 1998; McCown et al., 1996); however, innate inflammatory responses were observed (Lowenstein et al., 2007). Results reported by these methods are rooted in single-marker staining and thus prevent the discovery of unexpected cell-type-specific responses. A comprehensive understanding of the processes underlying viral vector or transgene-mediated responses is critical for further optimizing AAV gene delivery vectors and treatment modalities that mitigate such immune responses.

Here, we introduce an experimental and bioinformatics workflow capable of profiling the viral tropism and response of multiple barcoded AAV variants in a single animal across numerous complex cell types by taking advantage of the transcriptomic resolution of scRNA-seq techniques (Figure 1 A). For this proof-of-concept study, we profile the tropism of previously-characterized AAV variants that emerged from directed evolution with the CREATE (AAV-PHP.B, AAV-PHP.eB) (Chan et al., 2017; Deverman et al., 2016) or M-CREATE (AAV-PHP.C1, AAV-PHP.C2, AAV-PHP.V1, AAV.CAP-B10) (Flytzanis et al., 2020; Ravindra Kumar et al., 2020) platforms. We selected the AAV variants based on their unique CNS tropism following intravenous injection. AAV-PHP.B and AAV-PHP.eB are known to exhibit overall increased targeting of the CNS compared with AAV9 and preferential targeting of neurons and astrocytes. Despite its sequence similarity to AAV-PHP.B, the tropism of AAV-PHP.V1 is known to be biased toward transducing brain vascular cells. AAV-PHP.C1 and AAV-PHP.C2 have both demonstrated enhanced blood– brain barrier (BBB) crossing relative to AAV9 across two mouse strains (C57BL/6J and BALB/cJ). Finally, AAV.CAP-B10 is a recently-developed variant with a bias toward neurons compared to AAV-PHP.eB (Flytzanis et al., 2020).

**Figure 1.**
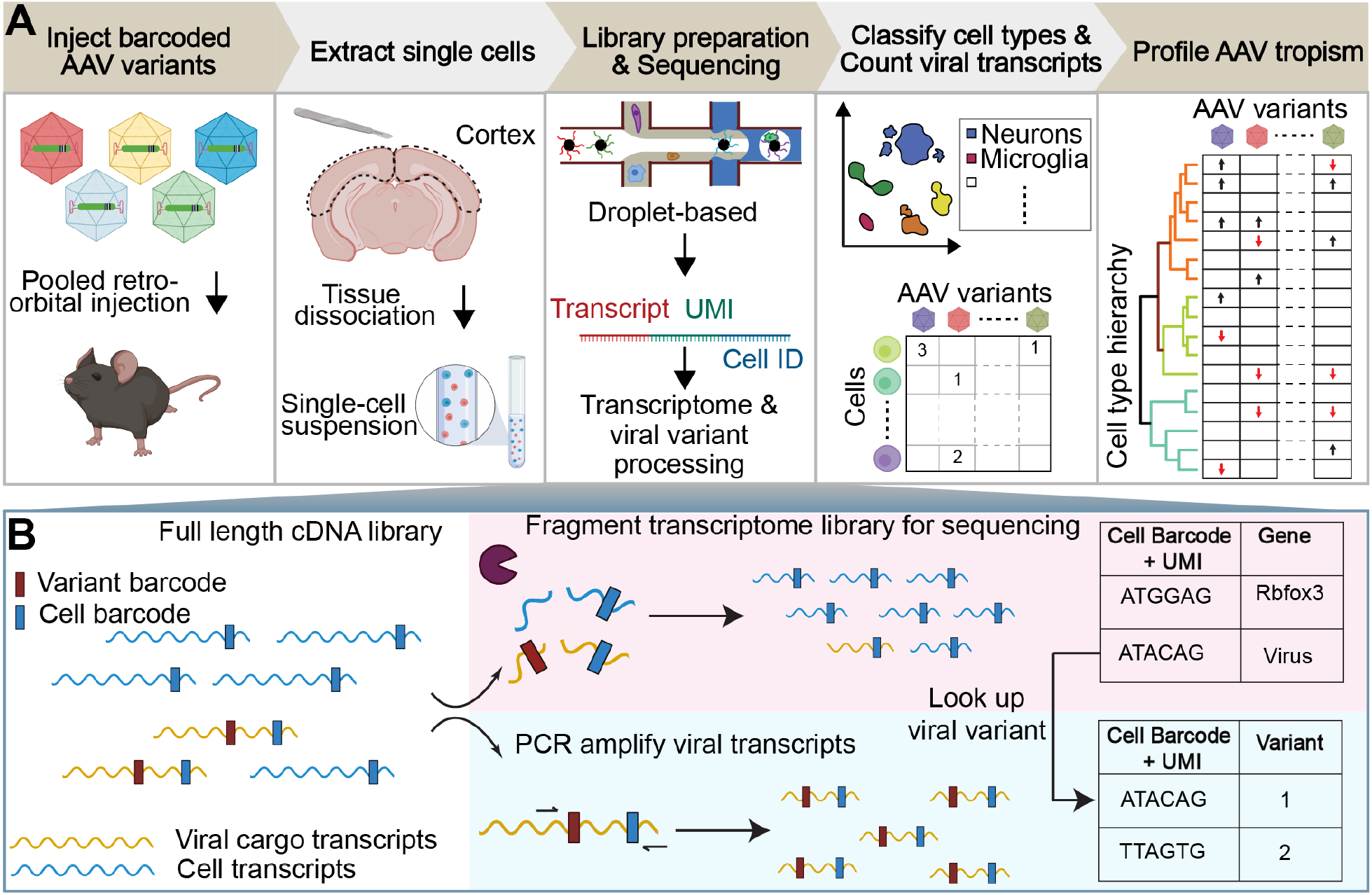
Workflow of AAV tropism characterization by scRNA-seq. (A) (I) Injection of a single AAV variant or multiple barcoded AAV variants into the retro-orbital sinus. (II) After 3–4 weeks post-injection, the brain region of interest is extracted and the tissue is dissociated into a single-cell suspension. (III) The droplet-based 10x Genomics Chromium system is used to isolate cells and build transcriptomic libraries (see B). (IV) Cells are assigned a cell-type annotation and a viral transcript count. (V) AAV tropism profiling across numerous cell types. (B) The full length cDNA library is fragmented for sequencing as part of the single-cell sequencing protocol (top). To enable viral tropism characterization of multiple rAAVs in parallel, an aliquot of the intact cDNA library undergoes further PCR amplification of viral transcripts (bottom). During cDNA amplification, Illumina sequencing primer targets are added to the viral transcripts such that the sequence in between the Illumina primer targets contains the AAV capsid barcode sequence. Viral cargo in the cell transcriptome is converted to variant barcodes by matching the corresponding cell barcode + UMI in the amplified viral transcript library (right).

In our initial validation experiment, we quantify the transduction biases of AAV-PHP.eB and AAV-CAP-B10 across major cell types using scRNA-seq, and our results correlate well with both published results and our own conventional IHC-based quantification. We then demonstrate the power of our transcriptomic approach by going beyond the major cell types to reveal significant differences in sub-cell-type transduction specificity. Compared with AAV-CAP-B10, AAV-PHP.eB displays biased targeting of inhibitory neurons, and both variants transduce Sst+ or Pvalb+ inhibitory neurons more efficiently than Vip+ inhibitory neurons. We validate these results with fluorescent in situ hybridization – hybridization chain reaction (FISH-HCR). We then develop and validate a barcoding strategy to investigate the tropism of AAV-PHP.V1 relative to AAV-PHP.eB in non-neuronal cells and reveal that pericytes, a subclass of vascular cells, evade transduction by this and other variants. We further use scRNA-seq to profile cell-type-specific responses to AAV.PHP-eB at 3 and 25 days post-injection (DPI), finding, for example, numerous genes implicated in the p53 pathway in endothelial cells to be upregulated at 3 DPI. While most upregulated genes across cell types return to control levels by day twenty-five, excitatory neurons show a persistent upregulation of genes involved in MAPK signaling extending to 25 days. Finally, we showcase the capabilities of parallel characterization by verifying the preceding findings in a single animal with seven co-injected AAV variants and reveal the unique non-neuronal tropism bias of AAV-PHP.C2.

## Results

### Multiplexed single-cell RNA sequencing-based AAV profiling pipeline

To address the current bottleneck in AAV tropism profiling, we devised an experimental and computational workflow (Figure 1 A) that exploits the transcriptomic resolution of scRNA-seq to profile the tropism of multiple AAV variants across complex cell-type hierarchies. In this workflow, single or multiple barcoded rAAVs are injected into the retro-orbital sinus of mice followed by tissue dissociation, single-cell library construction using the 10X Genomics Chromium system, and sequencing with multiplexed Illumina next-generation sequencing (NGS) (Zheng et al., 2017). The standard mRNA library construction procedure includes an enzymatic fragmentation step that truncates the cDNA amplicon such that its final size falls within the bounds of NGS platforms (Figure 1 B). These cDNA fragments are only approximately 450 bp in length and, due to the stochastic nature of the fragmentation, sequencing from their 5’ end does not consistently capture any particular region. The fragment length limit and heterogeneity pose a problem for parallelizing AAV tropism profiling, which requires reliable recovery of regions of the transgene that identify the originating AAV capsid. For example, posttranscriptional regulatory elements, such as the 600 bp Woodchuck hepatitis virus posttranscriptional regulatory element (WPRE), are commonly placed at the 3’ end of viral transgenes to modulate transgene expression. The insertion of such elements pushes any uniquely identifying cargo outside the 450 bp capture range, making them indistinguishable based on the cDNA library alone (Supplemental Figure 1 A). An alternative strategy of adding barcodes in the 3’ polyadenylation site also places the barcode too distant for a 5’ sequencing read, and reading from the 3’ end would require sequencing through the homopolymeric polyA tail, which is believed to be unreliable in NGS platforms (Chang et al., 2014; Shin and Park, 2016).

We circumvented these limitations in viral cargo identification by taking an aliquot of the intact cDNA library and adding standard Illumina sequencing primer recognition sites to the viral transcripts using PCR amplification such that the identifying region is within the two Illumina primer target sequences (e.g. Figure 2 B). The cell transcriptome aliquots undergoing the standard library construction protocol and the amplified viral transcripts are then sequenced as separate NGS libraries. We sequence shorter viral transcripts in the same flow cell as the cell transcriptomes and longer viral transcripts on the Illumina MiSeq, which we found to be successful at sequencing cDNAs up to 890 bp long. The sequencing data undergoes a comprehensive data processing pipeline (see Methods). Using a custom genome reference, reads from the cell transcriptome that align to the viral cargo plasmid sequences are counted as part of the standard 10X Cell Ranger count pipeline (see Methods and Supplemental Figure 1 C). In parallel, reads from the amplified viral transcripts are used to count the abundance of each viral barcode associated with each cell barcode and unique molecular identifier (UMI). The most abundant viral barcode for each cell barcode and UMI is assumed to be the correct viral barcode, and is used to construct a variant lookup table. This lookup table approach identifies an originating capsid in 67.6 ± 2.0% of viral transcripts detected in the cell transcriptome aliquots (Supplemental Table 4).

**Figure 2.**
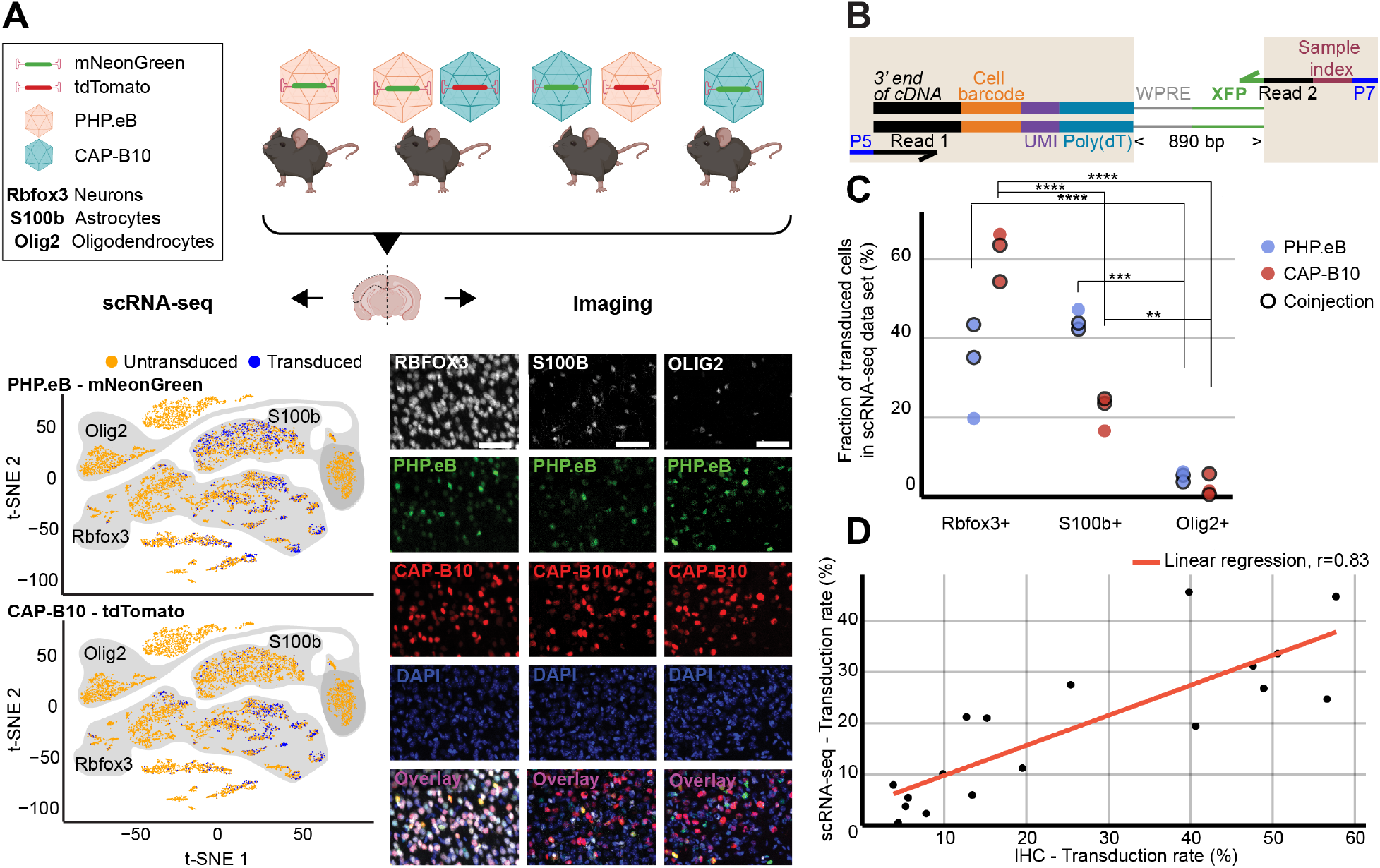
Comparison of viral tropism profiling with traditional IHC and scRNA-seq. (A) Overview of the experiment. Four animals were injected with 1.5 × 10^11^ viral genomes (vg) packaged in AAV-PHP.eB and/or AAV-CAP-B10. The bottom panels show a representative dataset collected from an animal that was co-injected with AAV-PHP.eB and AAV-CAP-B10. The left side displays the scRNA-seq data set in the lower dimensional t-SNE space, with cells colored according to transduction status. The shaded areas indicate clusters with high expression of the corresponding gene marker. The right side shows representative confocal images of cortical tissue labeled with IHC. Scale bar, 50 µm. (B) Viral transcript recovery strategy. The shaded areas highlight sequences added during library construction. (C) The fraction of the total number of transduced cells labeled as expressing the corresponding marker gene. For each AAV variant, the results of a two-way ANOVA with correction for multiple comparisons using Sidak’s test are reported with adjusted P-values (****P ≤ 0.0001, ***P ≤ 0.001, and **P ≤ 0.01 are shown; P > 0.05 is not shown). (D) Comparison of transduction rates based on quantification via scRNA-seq or IHC. Transduction rate was calculated as (number of transduced cells in the group)/(total number of cells in the group). Each dot represents the transduction rate of neurons/Rbfox3+, astrocytes/S100b+, or oligodendrocytes/Olig2+ by AAV-PHP.eB or AAV-CAP-B10 in one animal. Histology data are averages across three brain slices per gene marker and animal. r indicates the Pearson correlation coefficient.

For determining viral cell-type tropism, we developed a method to estimate the fraction of cells within a cell type that express viral transcripts. Viral RNA expression levels depend on both the multiplicity of infection and the transcription rate of the delivered cargo. Thus, directly using viral RNA counts to determine tropism is confounded by differences in transcription rate between cell types, limiting comparison with imaging-based tropism quantification methods. As evidence of this, we detected that viral RNA expression levels can vary by cell type but are not perfectly rank correlated with the percent of cells detected as expressing that transcript (Supplemental Figure 2 B). An additional confound arises from the ambient RNA from cellular debris co-encapsulated with cell-containing droplets, which can lead to false positives, i.e., detecting viral RNA in droplets containing a cell that was not expressing viral RNA. For example, we detected low levels of viral transcripts in large percentages of cells, even in cell types suspected to evade transduction, such as immune cells (Supplemental Figure 2 A). To reduce the effect of both variability in expression and ambient RNA, we developed an empirical method to estimate the percentage of cells expressing transcripts above the noise, wherein the distribution of viral transcript counts in a set of cells of interest is compared to a background distribution of cell-free (empty) droplets (see Methods, Supplemental Figure 2 C). In simulation, this method accurately recovers the estimated number of cells expressing transcripts above background across a wide range of parameterizations of negative binomial distributions (see Methods, Supplemental Figure 2 D).

To address several additional technical problems in default single-cell pipelines, we developed a simultaneous quality control (QC) and droplet identification pipeline. Our viral transduction rate estimation method described above relies on having an empirical background distribution of viral transcript counts in empty droplets to compare against the cell type of interest. However, the default cell vs. empty droplet identification method provided by the 10X Cell Ranger software, which is based on the EmptyDrops method (Lun et al., 2019), yielded unexpectedly high numbers of cells and clusters with no recognizable marker genes, suggesting they may consist of empty droplets of ambient RNA or cellular debris (Supplemental Figure 3 A, B). Additionally, we sought to remove droplets containing multiple cells (multiplets) from our data due to the risk of falsely attributing viral tropism of one cell type to another. However, using Scrublet (Wolock et al., 2019), an established method for identifying droplets containing multiplets, failed to identify multiplets in some of our samples and only identified small proportions of clusters positive for known non-overlapping marker genes, such as Cldn5 and Cx3cr1 (Supplemental Figure 3 C). To address both the empty droplet and multiplet detection issues, we built a droplet classification pipeline based on scANVI, a framework for classifying single-cell data via neural-network-based generative models (Xu et al., 2021). Using clusters with a high percentage of predicted multiplets from Scrublet as training examples of multiplets, and clusters positive for known neuronal and non-neuronal marker genes as training examples of neurons and non-neuronal cells, we trained a predictive model to classify each droplet as a neuron, non-neuron, multiplet, or empty droplet (see Methods, Supplemental Figure 4 A). This model performed with 97.6% accuracy on 10% of cells held out for testing, and yielded a database of 270,982 cortical cells (Supplemental Figure 4 B). Inspection of the cells classified as empty droplets reveals that these droplets have lower transcript counts and higher mitochondrial gene ratios, consistent with other single-cell quality control pipelines (Supplemental Figure 4 D). Critically, we discovered that non-neuronal clusters contained significantly more cells that had been previously removed by the Cell Ranger filtering method as compared to neuronal clusters (P = 0.02, 2-sided student t-test). In some clusters, such as Gpr17^+^ C1ql1^+^ oligodendrocytes and Gper^+^ Myl9^+^ vascular cells, we identified up to 85% more cells than what were recovered via Cell Ranger in some samples.

Using our combined experimental and computational pipeline for viral transcript recovery and droplet identification, we can recover a lower bound on the expected number of cells expressing each unique viral cargo within groups of cells in heterogeneous samples.

### Single-cell RNA sequencing recapitulates AAV capsid cell-type-specific tropisms

As a first step, we validated our method by comparing the quantification of AAV transduction of major cell types via scRNA-seq to conventional IHC. For this purpose, we characterized the tropism of two previously reported AAV variants, AAV-PHP.eB (Chan et al., 2017) and AAV-CAP-B10 (Flytzanis et al., 2020) (Figure 2 A). In total, four animals received single or dual retro-orbital injections of AAV-PHP.eB and/or AAV-CAP-B10 with 1.5 × 10^11^ viral genomes (vg) per variant. Co-injection of both variants served to test the ability of our approach to parallelize tropism profiling. By having each variant package a distinct fluorophore, tropism could be simultaneously assessed via multi-channel fluorescence and mRNA expression of the distinct transgene. After 3–4 weeks of expression, we harvested the brains and used one hemisphere for IHC and one hemisphere for scRNA-seq. To recover viral transcripts, we chose primers such that enough of the XFP sequence was contained within the Illumina primer target sequences to differentiate the two variants (Supplemental Table 1). For this comparison, we focused on the transduction rate for neurons (Rbfox3), astrocytes (S100b), and oligodendrocytes (Olig2). For IHC, a cell was classified as positive for the marker gene on the basis of antibody staining, and was classified as transduced on the basis of expression of the delivered fluorophore. For scRNA-seq, all cells that passed our QC pipeline were projected into a joint scVI latent space and clustered. To most closely match our imaging quantification, we considered all clusters that were determined to be positive for the respective marker gene as belonging to the corresponding cell type (see Methods). All clusters of the same marker gene were grouped together, and the transduction rate of the combined group of cells was determined using our viral transduction rate estimation method.

Our analysis of the scRNA-seq data demonstrates that the viral tropism biases across the three canonical marker genes are consistent with previous reports (Figure 2 C) (Chan et al., 2017; Flytzanis et al., 2020). In contrast to AAV-PHP.eB, AAV-CAP-B10 preferentially targets neurons over astrocytes and oligodendrocytes. No marked discrepancies in viral tropism characterization were observed with single versus dual injections.

To quantify the similarity of the AAV tropism characterizations obtained with IHC and scRNA-seq, we directly compared the transduction rate of each AAV variant for every cell type and its corresponding marker gene (i.e., Rbfox3, S100b, or Olig2) as determined by each technique and noticed a good correlation (Figure 2 D). Despite the different underlying biological readouts– protein expression in IHC and RNA molecules in labeled cell types for scRNA-seq–the two techniques reveal similar viral tropisms.

### Tropism profiling at transcriptomic resolution reveals AAV variant biases for neuronal subtypes

After validating our approach against the current standard of AAV tropism characterization (IHC imaging), we scrutinized the tropism of AAV-PHP.eB and AAV-CAP-B10 beyond the major cell types (Figure 3). Since AAV-CAP-B10 has increased neuronal bias relative to AAV-PHP.eB, we first sought to understand if there were neuronal subtypes that were differentially responsible for this bias. However, in-depth cell typing of transcriptomes collected from tissues with numerous and complex cell types, such as neurons in the brain, requires expert knowledge of the tissue composition, time to manually curate the data, and the availability of large datasets (Zeisel et al., 2018). To minimize the burden of manual annotation, computational tools have been developed that use previously-annotated single-cell databases to predict the cell type of cells in new, unannotated single-cell experiments, even across single-cell platforms (Cao et al., 2020; Tan and Cahan, 2019; Xu et al., 2021). We decided to leverage these tools and expanded our marker gene-based cell typing approach by having more complicated or well-established cell types be assigned based on annotations in a reference dataset (Supplemental Figure 4 A). To this end, we again employed scANVI to construct a joint model of cells from our samples and cells from an annotated reference database. For this model, we used the Mouse Whole Cortex and Hippocampus 10x v2 dataset available from the Allen Brain Institute (Yao et al., 2021). Since this is a neuron-enriched dataset, we constructed the model using only the 109,992 cells in our dataset classified as neurons from our marker-based QC pipeline combined with the 561,543 neuronal cells from cortical regions from the reference database. We trained this model to predict to which of 14 neuron subtype groupings each cell belonged. We held out 10% of the data for testing: the model performed with 97.9% classification accuracy on the held-out data. We then applied the model to predict the neuron subtypes of our cells.

**Figure 3.**
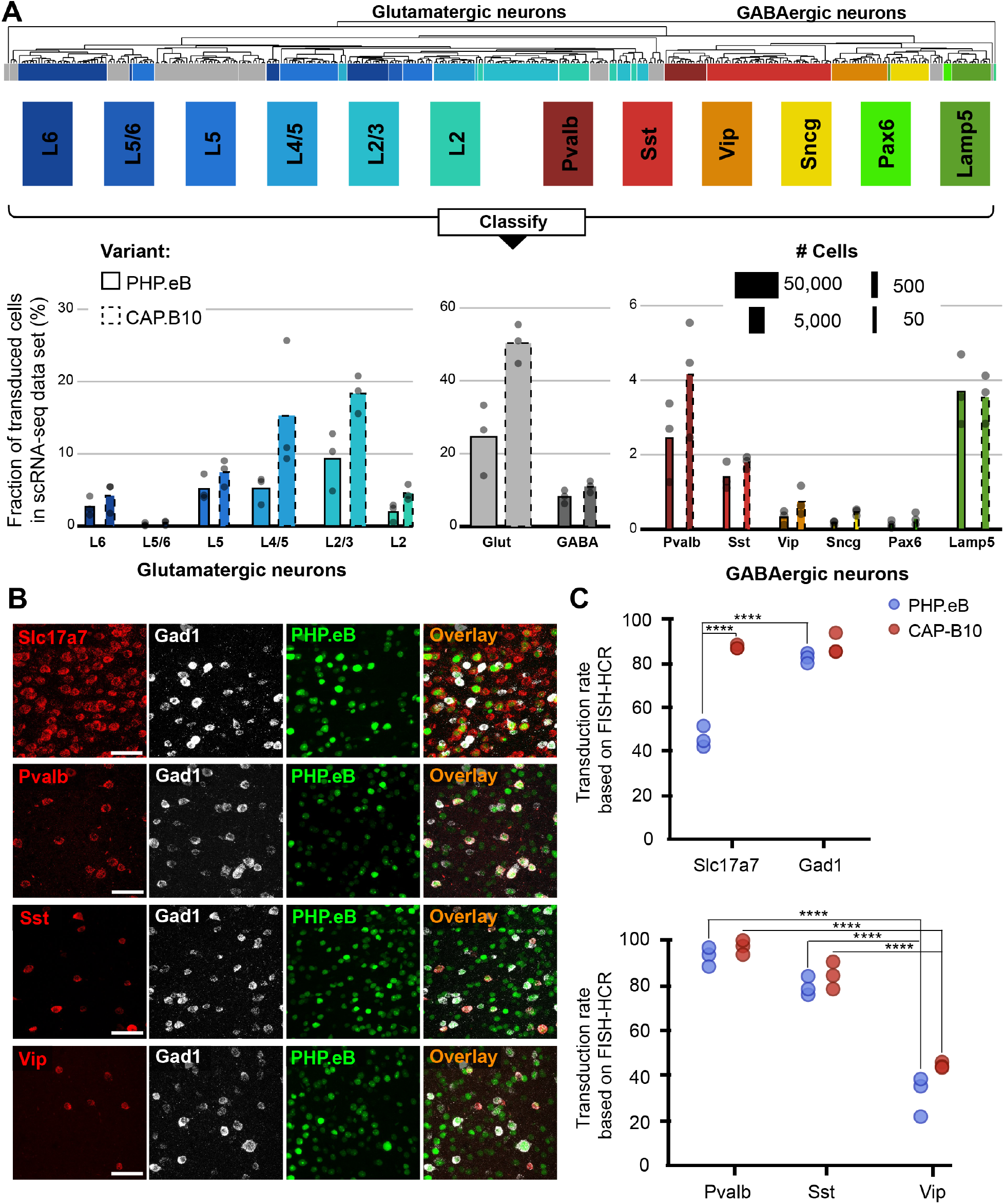
In-depth AAV tropism characterization of neuronal subtypes at transcriptomic resolution. (A) Viral tropism profiling across neuronal sub types. Neuronal subtype annotations are predicted by a model learned from the Allen Institute reference dataset using scANVI (Xu et al., 2021; Yao et al., 2021). Each dot represents data from one animal injected with AAV-PHP.eB and/or AAV-CAP-B10. Bar width indicates the total number of cells of a particular cell type present in our dataset. (B) Representative confocal images of cortical tissue from an animal injected with 1.5 × 10^11^ vg of AAV-PHP.eB. Tissue was labeled with FISH-HCR for gene markers of glutamatergic neurons (Slc17a7) and GABAergic neurons (Gad1, Pvalb, Sst, Vip). AAV-PHP.eB shows the endogenous fluorescence of mNeonGreen. Scale bar, 50 µm. (C) Confirmation of viral tropism biases across neuronal subtypes using FISH-HCR (3 mice per AAV variant, 1.5 × 10^11^ vg dose). Dots represent the average values across three brain slices from one animal. Results from a two-way ANOVA with correction for multiple comparisons using Tukey’s test is reported with adjusted P-values (****P ≤ 0.0001; and P > 0.05 is not shown on the plot).

During our in-depth characterization, we discovered several previously unnoticed sub-cell-type biases for AAV-PHP.eB and AAV-CAP-B10 (Figure 3 A). Starting at the top of our neuronal hierarchy, the fraction of transduced cells that were glutamatergic neurons was markedly reduced for AAV-PHP.eB compared with AAV-CAP-B10 (P = 0.03, 2-sided student t-test, corrected for 2 neuron subtype comparisons). Furthermore, Pvalb+ and Sst+ inhibitory neurons both represented a larger fraction of transduced cells than Vip+ inhibitory neurons with both variants (adjusted P < 0.0001, P = 0.10, respectively, two-way ANOVA with multiple comparison correction for inhibitory neuron subtypes using Tukey’s method).

To confirm these tropism biases in neuronal subtypes with a traditional technique, we performed FISH-HCR for glutamatergic and GABAergic gene markers (Figure 3 B) (Choi et al., 2014; Patriarchi et al., 2018). As indicated by our scRNA-seq data, AAV-CAP-B10, when compared with AAV-PHP.eB, has increased transduction efficiency of glutamatergic neurons (SLC17A7). Furthermore, FISH-HCR verified the downward trend in transduction efficiency from Pvalb+, to Sst+, to Vip+ neurons in both AAV variants (Figure 3 C).

### Pooled AAVs packaging barcoded cargo recapitulate the non-neuronal tropism bias of PHP.V1

To enable profiling viral variants in parallel without needing distinct transgenes per variant, we established a barcoding strategy whereby we package AAV variants with the same transgene and regulatory elements but with short, distinguishing nucleotide sequences within the 3’ UTR (Figure 4 A). To verify that this barcoding strategy can recover tropisms consistent with our previous transgene-based capsid-identification strategy, we performed a set of experiments to re-characterize the tropism of AAV-PHP.eB in parallel with that of the recently developed AAV-PHP.V1, which has increased specificity for vascular cells over AAV-PHP.eB (Ravindra Kumar et al., 2020).

**Figure 4.**
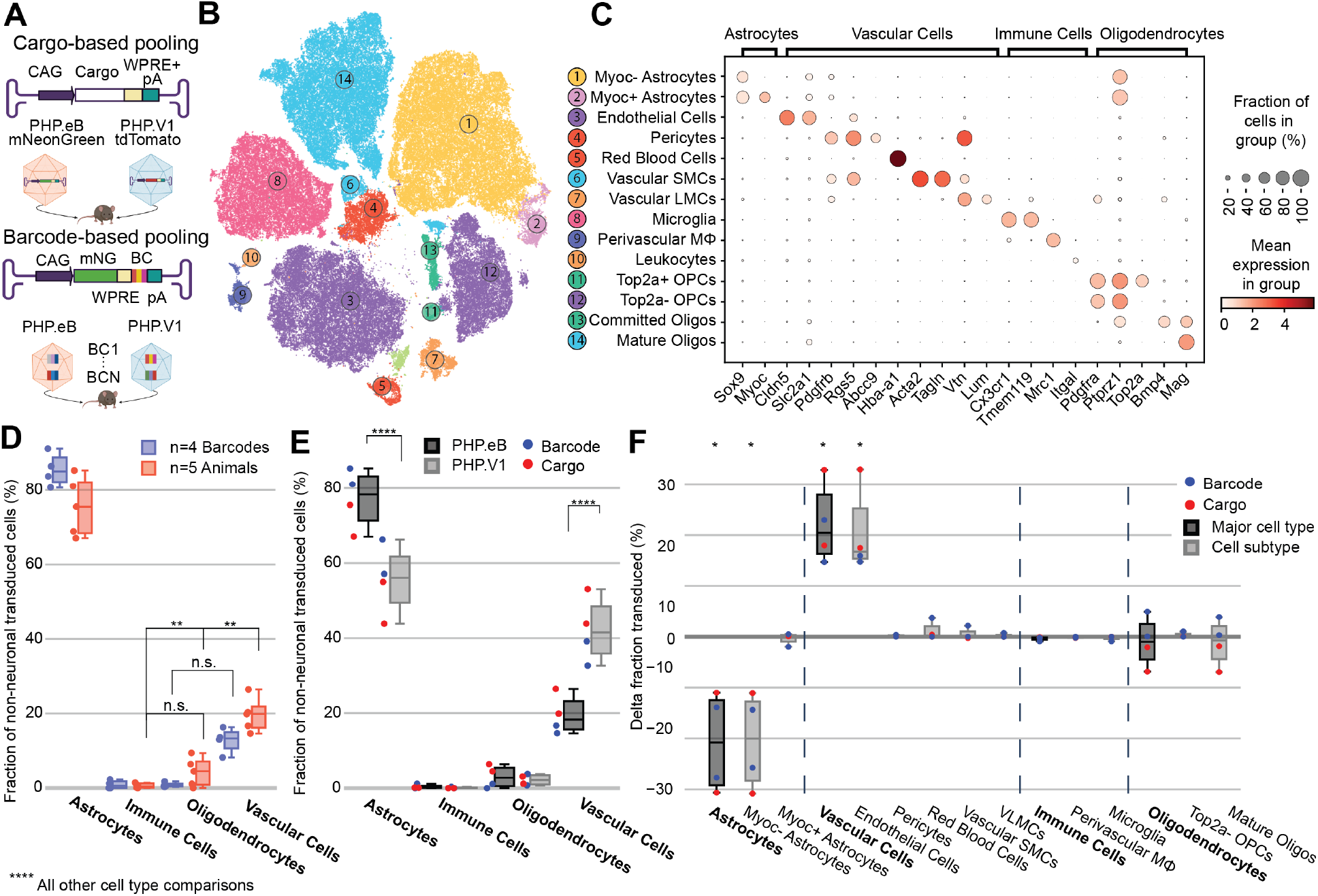
Barcoded co-injected rAAVs reveal the non-neuronal tropism bias of AAV-PHP.V1. (A) Experimental design for comparing barcode vs cargo-based tropism profiling. Animals received dual injections of AAV-PHP.eB and AAV-PHP.V1, carrying either distinct fluorophores (cargo) or the same fluorophore with distinct barcodes. (B) t-SNE projection of the single-cell Variational Inference (scVI) latent space of cells and their cell type classification of the 169,265 non-neuronal cells across all our samples. Each number corresponds to the cell type labeled in (C). (C) Marker genes used to identify non-neuronal cell types. Darker colors indicate higher mean expression, and dot size correlates with the abundance of the gene in that cell type. (D) The distribution of non-neuronal cells expressing transcripts from AAV-PHP.eB across 4 barcodes within one animal (blue) and across 5 animals (red). All animals received dual injections, with one of the vectors being 1.5 x 10^11^ vg of PHP.eB carrying CAG-mNeonGreen. The y-axis represents the fraction of transduced non-neuronal cells that are of the specified cell type. Only the non-significant comparisons between cell types in a two-way ANOVA with correction for multiple comparisons using Tukey’s test are reported. All other cell-type comparisons within a paradigm were significant at P ≤ 0.0001. (E) The distribution of non-neuronal cells expressing transcripts from AAV-PHP.eB (black) and AAV-PHP.V1 (gray). Results from the different experimental paradigms are combined. Results shown are from a two-way ANOVA with correction for multiple comparisons using Sidak’s test comparing transduction by AAV-PHP.eB to AAV-PHP.V1 for each cell type, with adjusted P-values (****P ≤ 0.0001 is shown; P > 0.05 is not shown). (F) Within-animal difference in the fraction of cells transduced with AAV-PHP.V1 relative to AAV-PHP.eB across four animals, two from each experimental paradigm. For each cell type in each sample, the combined 2-proportion z score for the proportion of that cell type transduced by AAV-PHP.V1 vs AAV-PHP.eB is reported. Cell types with fewer than 2 cells transduced by both variants were discarded. Z scores were combined across multiple animals using Stouffer’s method and corrected for multiple comparisons. Cell-type differences with an adjusted P-value below 0.05 are indicated with *.

We produced AAV-PHP.eB carrying CAG-mNeonGreen and AAV-PHP.V1 carrying either CAG-mRuby2 or CAG-tdTomato. Additionally, we produced AAV-PHP.eB and AAV-PHP.V1 both carrying CAG-mNeonGreen with 7-nucleotide barcodes 89 bp upstream of the polyadenylation start site such that they did not interfere with the WPRE. We ensured each barcode had equal G/C content, and that all barcodes were Hamming distance 3 from each other (Supplemental Table 5). Each of the barcoded variants was packaged with multiple barcodes that were pooled together during virus production. Four animals received a retro-orbital co-injection of 1.5 x 10^11^ vg/each of AAV-PHP.V1 and AAV-PHP.eB. Two animals received viruses carrying separate fluorophores (cargo-based), and two animals received viruses carrying the barcoded cargo (barcode-based). For amplification of the viral cDNA in the animals receiving the barcoded cargo, we used primers closer to the polyA region such that the sequencing read covered the barcoded region (Supplemental Table 1). During the single-cell sequencing dissociation and recovery, one of our dissociations resulted in low recovery of neurons (Supplemental Figure 4 C); thus, we investigated only non-neuronal cells for this experiment.

Despite variability in the total transgene RNA content between barcodes of the same variant (Supplemental Figure 5 A), the estimated percent of cells expressing the transgene within each cell type was consistent between barcodes within a single animal, with standard deviations ranging from 0.003 to 0.058 (Supplemental Figure 6 A). Our analysis of both the barcode-based animals and cargo-based animals shows the same bias in non-neuronal tropism, with AAV-PHP.eB significantly preferring astrocytes over oligodendrocytes, vascular cells, and immune cells (Figure 4 D). Interestingly, our analysis also revealed that the variance between barcodes within an animal was less than the variance between animals, even when controlling for cargo and dosage (P = 0.021, Bartlett’s test, P-values combined across all variants and cell types using Stouffer’s method, weighted by transduced cell type distribution).

Next, we investigated the distribution of cells transduced by AAV-PHP.eB vs AAV-PHP.V1 in the major non-neuronal cell types across both barcode-based and cargo-based paradigms (Figure 4 E). The single-cell tropism data confirms the previously-established finding that AAV-PHP.V1 has a bias toward vascular cells relative to AAV-PHP.eB. Additionally, we uncovered that this is coupled with a bias away from astrocytes relative to AAV-PHP.eB, but that transduction of oligodendrocytes and immune cells did not differ between the variants. To investigate for a specific effect of the barcoding strategy, we performed a three-way ANOVA across the variant, cell type, and experimental paradigm factors. We found that the cell type factor accounted for 89.25% of the total variation, the combined cell type + variant factor accounted for 7.7% of the total variation, and the combined cell type + experimental paradigm factor accounted for only 2.0% of the total variation, confirming our hypothesis that barcoded pools can recover tropism with minimal effect.

### Relative tropism biases reveal non-neuronal subtypes with reduced AAV transduction

To further characterize the tropism biases of AAV-PHP.V1 and expand our method to less well-established cell hierarchies, we explored the non-neuronal cell types in our dataset. Since the Allen Brain Institute reference database that we used to investigate neuronal tropism was enriched for neurons, it does not contain enough non-neuronal cells to form a robust non-neuronal cell atlas. Our combined dataset consists of 169,265 non-neuronal cells, making it large enough to establish our own non-neuronal cell clustering. Thus, we performed an additional round of automatic clustering on the cells classified as non-neuronal in our combined dataset, and identified 12 non-neuronal cell subtypes based on previously established marker genes (Figure 4 B, C, Supplemental Table 2).

Most cell subtypes had multiple clusters assigned to them, which suggested there may be additional subtypes of cells for which we did not find established marker genes. To determine whether any of these clusters delineated cell types with distinct transcriptional profiles, we investigated the probability of gene expression in each cluster compared to the other clusters of the same cell subtype (see Methods). Our approach determined two subclusters of pericytes, astrocytes, and oligodendrocyte precursor cells (OPCs). Both clusters of pericytes had strong expression of canonical pericytes marker genes Rgs5, Abcc9, and Higd1b. However, one of the clusters had no marker genes that made it distinct from the other pericyte cluster, nor from endothelial cells. Consistent with previous reports, this suggests that this cluster could be pericytes contaminated with endothelial cell fragments, and thus was not considered for further analysis (He et al., 2016; Vanlandewijck et al., 2018; Yang et al., 2021). Two distinct groups of astrocytes were detected, one of which had unique expression of Myoc and Fxyd6. Finally, one of the clusters of OPCs were uniquely expressing Top2a, Pbk, Spc24, Smc2, and Lmnb1. Using these new marker genes, we expanded our non-neuronal cell taxonomy to 14 cell types, now including Myoc+ and Myoc-astrocytes, and Top2a+ and Top2a-OPCs.

Given our finding that inter-sample variability exceeds intra-sample variability, we established a normalization method for comparing transduction biases between variants co-injected into the same animal. This normalization–calculating the difference in the fraction of transduced cells between variants–captures the relative bias between variants, instead of the absolute tropism of a single variant (see Methods). By considering the relative bias between variants, we are able to interrogate tropism in a way that is more robust to inter-sample variability that arises from different distributions of recovered cells, expression rate of delivered cargo, and success of the injection. Using this normalization method, we evaluated the non-neuronal cell type bias of AAV-PHP.V1 relative to AAV-PHP.eB in both the cargo-based animals and the barcode-based animals across our non-neuronal cell-type taxonomy (Figure 4 F). We discovered that the bias of AAV-PHP.V1 for vascular cells is driven by an increase in transduction of endothelial cells, but not pericytes.

Similarly, AAV-PHP.V1’s bias away from astrocytes is driven by a decrease in transduction of Myoc-astrocytes, but not Myoc+ astrocytes. Further inspection of the transduction of pericytes and Myoc+ astrocytes revealed that pericytes are not highly transduced by any of the AAVs tested in this work, and that Myoc+ astrocytes have both lower viral transcript expression and lower abundance than Myoc-astrocytes, and thus do not contribute significantly to tropism (Supplemental Figure 4, 7 A, B).

### Single-cell RNA sequencing reveals early cell-type-specific responses to IV administration of AAV-PHP.eB that return to baseline by 3.5 weeks

To investigate the temporal cell-type-specific transcriptional effects of systemic AAV delivery and cargo expression, we performed a single-cell profiling experiment comparing animals injected with AAV to saline controls. We injected four male mice with AAV-PHP.eB (1.5 x 10^11^ vg) carrying mNeonGreen, and performed single-cell sequencing on two mice three days post-injection (3 DPI) and two mice twenty-five days post-injection (25 DPI). These time points were chosen based on previous work showing MHC presentation response peaking around day seven and transgene response peaking around day 30 (Lowenstein et al., 2007). The two saline control mice were processed 3 DPI. We then analyzed differential gene expression for each cell type between injected animals and controls using DESeq2 (Supplemental Table 7). Of note, we excluded cell types with less than 50 cells in each sample, and excluded leukocytes and red blood cells given the risk of their presence due to dissociation rather than chemokine mediated infiltration. Additionally, we collapsed subtypes of excitatory neurons, inhibitory neurons, and OPCs to have greater than 50 cells for differential analysis. We estimated viral transduction rate of AAV-PHP.eB using its delivered cargo, mNeonGreen, across cell types and time points. We identified that Myoc-Astrocytes have significantly higher estimated transduction rate at 25 DPI compared to 3DPI (adjusted P-value = 0.0003, two-way ANOVA with multiple comparison correction using Sidak’s method). It is also worth noting that endothelial cells have a similar transduction rate between the time points in both animals, while one of the animals at 25 DPI exhibited higher transduction in neurons (Figure 5A). The number of statistically relevant genes between the injected and control group (adjusted P-value < 0.05, DESeq2) were highest in pericytes (26 genes), endothelial cells (76 genes), and excitatory neurons (45 genes) at 3 DPI (Figure 5 B). At day twenty-five, only excitatory neurons had greater than 10 genes (14 genes total) differentially expressed (adjusted P-value < 0.05, DESeq2).

**Figure 5.**
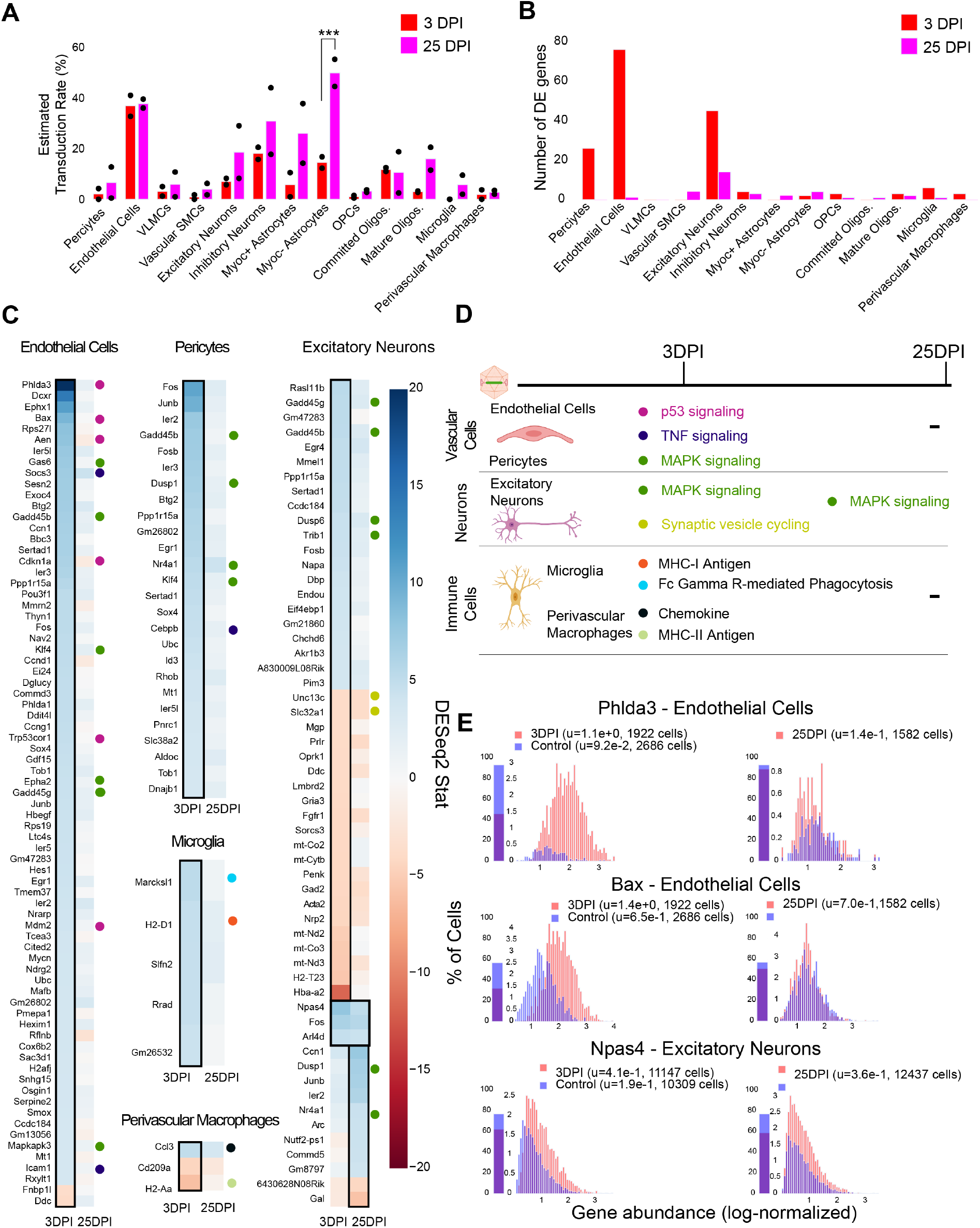
Single-cell gene expression profiling finds cell-type-specific responses to AAV transduction in vascular cells and excitatory neurons. (A) Estimated transduction rate (%) of mNeonGreen cargo at three and twenty-five days post-injection (DPI). Results from a two-way ANOVA with correction for multiple comparisons using Sidak’s method is reported with adjusted P-values (***P ≤ 0.001; and P > 0.05 is not shown on the plot). (B) Number of differentially expressed genes (adjusted P-value < 0.05, DESeq2) at 3 DPI and 25 DPI across 2 animals. (C) Differentially expressed genes across the two time points in endothelial cells, pericytes, microglia, perivascular macrophages, and excitatory neurons. Color indicates DESeq2 test statistic with red representing downregulation and blue representing upregulation. Genes outlined by a black rectangle are determined to have statistically significant differential expression compared to controls (adjusted P-value < 0.05, DESeq2). Colored circles adjacent to each gene indicate the corresponding pathway presented in (D). (D) A summary of corresponding pathways in which the differentially regulated genes in (C) are involved across the time points. (E) Distribution of p53 signaling transcripts in endothelial cells (animals are combined) and an example of a gene upregulated in both 3 and 25 DPI in excitatory neurons.

We found that endothelial cells had the most acute response at 3 DPI with pathways such as p53, MAPK, and TNF signaling notably impacted. A significant upregulation of Phlda3 and its effectors Bax, Aen, Mdm2, and Cdkn1a, all involved in the p53/Akt signaling pathway, was present (Figure 5 C, E) (Ferreira and Nagai, 2019; Ghouzzi et al., 2016). Of relevance, we also detected Trp53cor1/LincRNA-p21, responsible for negative regulation of gene expression (Amirinejad et al., 2020), upregulated in endothelial cells at 3 DPI. Other examples of upregulated genes relevant to inflammation and stress response in vascular cells include the suppressor of cytokine signaling protein Socs3 (Baker et al., 2009), and Mmrn2, responsible for regulating angiogenesis in endothelial cells (Lorenzon et al., 2012). Expression of Socs3 and Icam1, which are upregulated in endothelial cells at 3 DPI, and Cepbp, which is upregulated in pericytes at 3 DPI, have all been linked to TNF signaling (Burger et al., 1997; Cao et al., 2018; Li et al., 2020). We have also observed genes linked to MAPK signaling upregulated in endothelial cells, such as Gas6, Epha2, and Mapkapk3, and Klf4 in both endothelial cells and pericytes (Chen et al., 1997; Macrae et al., 2005; Riverso et al., 2017).

In brain immune cells, we observe a few substantial changes in genes pertaining to immune regulation at 3DPI which vanish at 25 DPI. For example, we observe an upregulation of MHC-I gene H2-D1 at 3 DPI in microglia, which then stabilizes back to control levels at 25 DPI (Figure 5 C). Marcksl1, previously reported as a gene marker for neuroinflammation induced by alpha-synuclein (Sarkar et al., 2020), also shows upregulation at 3 DPI. We did not observe significant differences in pro-inflammatory chemokines, Ccl2 and Ccl5, which are related to breakdown of the blood-brain barrier via regulation of tight-junction proteins and recruitment of peripheral leukocytes (Gralinski et al., 2009). Ccl3, responsible for infiltration of leukocytes and CNS inflammation (Chui and Dorovini-Zis, 2010), was upregulated in perivascular macrophages in 3 DPI and diminished back to control levels at 25 DPI (Figure 5 C). In contrast, Cd209a, a gene previously identified as critical for attracting and activating naïve T Cells (Franchini et al., 2019), was downregulated at 3 DPI.

Interestingly, we found that excitatory neurons had changes in genes across both 3 DPI and 25 DPI. MHC-Ib H2-T23, which is involved in the suppression of CD4+ T cell responses (Ohtsuka et al., 2008), is downregulated at 3 DPI. Meanwhile, the growth arrest genes, Gadd45g and Gadd45b (Vairapandi et al., 2002), are upregulated. Genes involved in synaptic vesicle cycling, such as Unc13c (Palfreyman and Jorgensen, 2017) and Slc32a1 (Taoufiq et al., 2020), are also downregulated at 3 DPI. Some genes remain upregulated throughout the study, such as Npas4, responsible for regulating excitatory-inhibitory balance (Spiegel et al., 2014). Genes implicated in MAPK signaling were upregulated – such as Gadd45b/g, Dusp6, and Trib1 at 3 DPI, and Dusp1 and Nr4a1 at 25 DPI (Muhammad et al., 2018; Ollila et al., 2012; Pérez-Sen et al., 2019; Salvador et al., 2013; Zhang and Yu, 2018). Gadd45b, Dusp1, Nr4a1 were also upregulated in pericytes and Gadd45b/g in endothelial cells (Figure 5 C).

Immediate early genes such as Ier2 (Kodali et al., 2020) were upregulated across pericytes, endothelial cells, inhibitory neurons, and OPCs at 3 DPI, while Fos, Ier2, Junb, and Arc were prominent in excitatory neurons at 25 DPI.

By investigating the gene expression differences in subpopulations of cells post-injection, we found that vascular cells such as endothelial cells and pericytes upregulate genes linked to p53, MAPK, and TNF signaling pathways at 3 DPI (Figure 5 D). Immune cells such as microglia and perivascular macrophages upregulate genes involved in chemokine signaling, MHCI/II antigen processing, and Fc Gamma R-Mediated Phagocytosis (Zhang et al., 2021) at 3 DPI (Figure 5 D). Excitatory neurons are the only cell type with genes implicated in the same pathway (MAPK signaling) upregulated across both of the time points (3 DPI, 25 DPI).

### Larger pools of barcoded AAVs recapitulate complex tropism within a single animal

To showcase the capabilities of parallel characterization, we next designed a 7-variant barcoded pool that included the three previously characterized variants (AAV-PHP.eB, AAV-CAP-B10, and AAV-PHP.V1), AAV9 and AAV-PHP.B controls, and two additional variants, AAV-PHP.C1 and AAV-PHP.C2. For simplification of cloning and virus production, we designed a plasmid, UBC-mCherry-AAV-cap-in-cis, that contained both the barcoded cargo, UBC-mCherry, and the AAV9 capsid DNA (Supplemental Figure 1 B). We assigned three distinct 24 bp barcodes to each variant (Supplemental Table 5). Each virus was produced separately to control the dosage, and 1.5 x 10^11^ vg of each variant was pooled and injected into a single animal.

After 3 weeks of expression, we performed single-cell sequencing on extracted cortical tissue. To increase the number of cells available for profiling, we processed two aliquots of cells, for a total of 36,413 recovered cells. To amplify the viral transcripts, we used primers that bind near the 3’ end of mCherry such that the barcode was captured in sequencing (Supplemental Table 1).

Using our cell typing and viral transcript counting methods, we investigated the transcript counts and transduction bias of the variants in the pool. Compared with our previous profiling experiments, the number of UBC-mCherry viral transcripts detected per cell was significantly lower than CAG-mNeonGreen-WPRE and CAG-tdTomato (adjusted P < 0.0001, P=0.0445, respectively, two-way ANOVA with multiple comparison correction using Tukey’s method) and shifted towards vascular cells (adjusted P < 0.0001, P=0.0008, respectively, two-way ANOVA with multiple comparison correction using Tukey’s method) (Supplemental Figure 5 B, C). Next, we looked at the transduction rate difference for each variant compared with the rest of the variants in the pool for each cell type in our taxonomy (Figure 6 A, B). Despite the lower expression rate and bias shift, the transduction rate difference metric captured the same tropism biases for AAV-CAP-B10 and AAV-PHP.V1 as determined from our previous experiments. AAV-CAP-B10 showed enhanced neuronal targeting relative to other variants in the pool, with this bias coming specifically from an increase in the transduction of glutamatergic neurons. All five variants with transcripts detected in neurons showed a decreased transduction rate in Vip+ neurons relative to other GABAergic neuronal subtypes (Supplemental Figure 7 C). AAV-PHP.eB showed enhanced targeting of astrocytes (+6.2%, P = 1.4 x 10^-8^, 2-proportion z-test, multiple comparison corrected with Benjamini/Hochberg correction), and AAV-PHP.V1 showed strong bias for vascular cells (+51.6%, p = 1.7 x 10^-43^). In addition to confirming all our existing hypotheses, we were able to identify biases for the previously reported AAV-PHP.C2, which has not been characterized in depth. This variant, which was reported as having a non-neuronal bias similar to AAV-PHP.V1, showed significant transduction bias not only toward vascular cells (+13.6%, P = 8.3 x 10^-6^), but also toward astrocytes (+24.0%, P = 1.6^-30^), and a bias away from neurons (−38%, p = 4.1 x 10^-32^).

**Figure 6.**
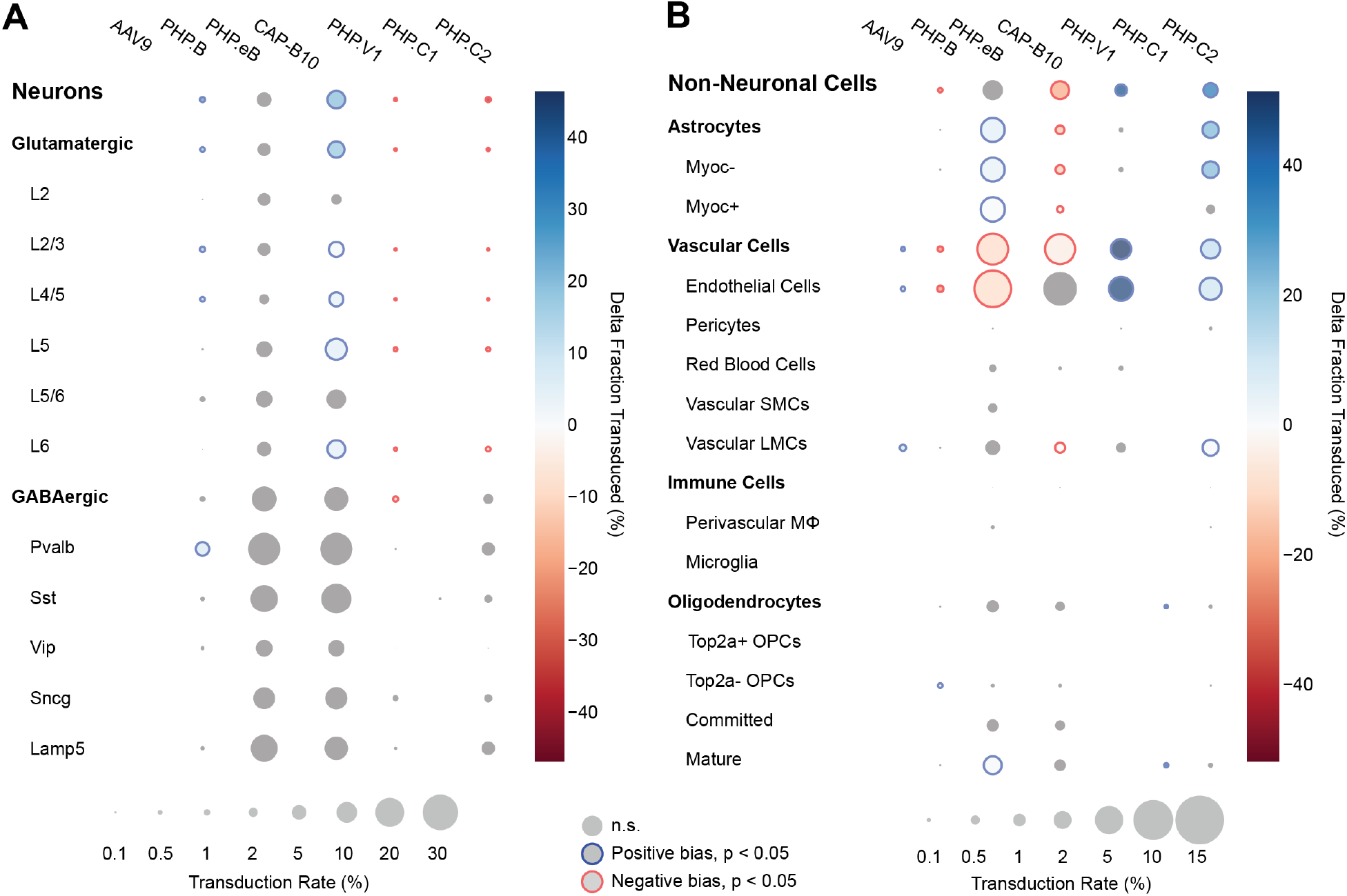
Single animal injections of multiple barcoded rAAVs enables deep, parallel characterization. (A, B) Relative cell type tropism of 7 co-injected rAAVs for neuronal (A) and non-neuronal (B) cell types. The color scale indicates the difference in transduction bias of a variant relative to all other variants in the pool. The area of each circle scales linearly with the fraction of cells of that type with viral transcripts above background. For each variant and cell type, a 2-proportion z score was calculated to compare the number of cells of that type transduced by that variant relative to all other variants combined. Z scores were combined across two single-cell sequencing aliquots using Stouffer’s method, and corrected for multiple comparisons. Cell types with fewer than 10 transduced cells in either the variant or variants compared against were discarded. Only cell-type biases at an adjusted P-value < 0.05 are colored; otherwise they are grayed out.

## Discussion

The advent of NGS has enabled screening of large libraries of AAV capsids in vivo by extracting viral DNA from relevant tissue followed by sequencing of capsid gene inserts or DNA barcodes corresponding to defined capsids. To date, NGS-based screening has been successfully applied to libraries created by peptide insertions (Davidsson et al., 2019; Körbelin et al., 2016b), DNA shuffling of capsids (De Alencastro et al., 2020; Herrmann et al., 2019; Paulk et al., 2018), and site-directed mutagenesis (Adachi et al., 2014). Although these NGS-based strategies allow the evolution of new AAV variants with diverse tissue tropisms, it has been difficult to obtain a comprehensive profiling for multiple variants across cell types, which is of utmost importance in organs with complex cell-type compositions, such as the brain (Deverman et al., 2016; Ravindra Kumar et al., 2020; Tasic et al., 2016, 2018; Zeisel et al., 2018). Towards this end, techniques such as IHC, fluorescent in situ RNA hybridization (Chen et al., 2015; Choi et al., 2014; Femino et al., 1998; Lubeck et al., 2014; Shah et al., 2016a, 2016b) or in situ RNA sequencing (Ke et al., 2013; Lee et al., 2014; Wang et al., 2018) can be employed. Several limitations make it challenging to apply these techniques as high-throughput, post-selection AAV tropism profiling methods. First, the limits of optical resolution and the density of transcripts in single cells pose challenges for full in situ transcriptome analysis and, until recently, have restricted the total number of simultaneously measured genes in single cells within tissue to several hundred (Ke et al., 2013; Lee et al., 2014; Liao et al., 2020; Shah et al., 2016a; Wang et al., 2018). By contrast, scRNA-seq with the 10x Genomics Chromium system enables detection of over 4000 genes per cell (Yao et al., 2021), fast transcriptomic analysis, and multiplexing across different tissue types (McGinnis et al., 2019; Stoeckius et al., 2018). Furthermore, the method is already widely used by the research community which can help with adoption of our proposed pipelines. Although droplet-based scRNA-seq methods lose spatial information during the dissociation procedure, analysis packages have been developed that can infer single-cell localization by combining scRNA-seq data with pre-existing information from ISH-based labeling for specific marker genes (Achim et al., 2015; Durruthy-Durruthy et al., 2015; Halpern et al., 2017; Nitzan et al., 2019; Satija et al., 2015; Stuart et al., 2019). Therefore, scRNA-seq techniques have great potential to rapidly profile the tropism of multiple AAV variants in parallel across several thousand cells defined by their entire transcriptome.

Here, we established an experimental and data-analysis pipeline that leverages the capabilities of scRNA-seq to achieve simultaneous characterization of several AAV variants across multiplexed tissue cell types within a single animal. To differentiate multiple AAV capsid variants in the sequencing data, we packaged variants with unique transgenes or the same transgene with unique barcodes incorporated at the 3’ end. We added standard Illumina sequencing primer recognition sites (Read 2) to the viral transcripts using PCR amplification such that the barcoded region could be consistently read out from the Illumina sequencing data. Our computational pipeline demultiplexes viral reads found in the transcriptome according to which matching sequence is most abundant in a separate amplified viral transgene library. Comparing the distribution of viral transcripts by cell type to a null model of empty droplets, we could then determine the cell-type biases.

Our platform has corroborated the tropism of several previously characterized AAV variants and has provided more detailed tropism information beyond the major cell types. The fraction of transduced cells that are glutamatergic neurons was found to be markedly reduced for AAV-PHP.eB when compared with AAV-CAP-B10. Furthermore, within all the variants we tested, both Pvalb+ and Sst+ inhibitory neurons have greater transduction rates than Vip+ neurons.

This bodes well for delivery to Pvalb+ neurons, which have been implicated in a wide range of neuro-psychiatric disorders (Ruden et al., 2021), and suggests Vip+ interneurons, which have recently been identified as being a sufficient delivery target for induction of Rett syndrome-like symptoms, as a target for optimization (Mossner et al., 2020). Awareness of neuronal subtype biases in delivery vectors is critical both for neuroscience researchers and for clinical applications. Dissection of neural circuit function requires understanding the roles of neuronal subtypes in behavior and disease and relies on successful and sometimes specific delivery of transgenes to the neuronal types under study (Bedbrook et al., 2018).

We further discovered that the vascular bias of AAV-PHP.V1 originates from its transduction bias towards endothelial cells. Interestingly, this is the only cell type we detected expressing Ly6a (Supplemental Figure 8), a known surface receptor for AAV variants in the PHP.B family (Batista et al., 2020; Hordeaux et al., 2019; Huang et al., 2019). Given AAV-PHP.V1’s sequence similarity to AAV-PHP.B and its tropism across mouse strains, this pattern suggests that AAV-PHP.V1 transduction may also be Ly6a-mediated. Finding such associations between viral tropism and cell-surface membrane proteins also suggests that full transcriptome sequencing data may hold a treasure trove of information on possible mechanisms of transduction of viral vectors.

We also revealed that AAV-PHP.C2 has a strong, broad non-neuronal bias toward both vascular cells and astrocytes. AAV-PHP.C2 also transduces BALB/cJ mice, which do not contain the Ly6a variant that mediates transduction by PHP.B family variants (Hordeaux et al., 2019). This suggests that PHP.C2 may be the most promising candidate from this pool for researchers interested in delivery to non-neuronal cells with minimal neuronal transduction both in C57BL/6J mice and in strains and organisms that do not have the Ly6a variant.

All our tested variants with non-neuronal transduction have lower expression in Myoc+ astrocytes and pericytes. Astrocytes expressing Myoc and Gfap, which intersect in our data (Supplemental Figure 8), have been previously identified as having reactive behavior in disease contexts, making them a target of interest for research on neurological diseases (Perez-Nievas and Serrano-Pozo, 2018; Wu et al., 2017). Similarly, pericytes, whose dysfunction has been shown to contribute to multiple neurological diseases, may be an important therapeutic target (Blanchard et al., 2020; Liu et al., 2020; Montagne et al., 2020). Both of these cell types may be good candidates for further AAV optimization, but may have been missed with marker gene-based approaches. In both AAV characterization and neuroscience research efforts, different marker genes are often used for astrocyte classification – sometimes more restrictive genes such as Gfap, and other times more broadly expressing genes such as S100b or Aldh1l1 (Yang et al., 2011; Zhang et al., 2019). Similarly, defining marker genes for pericytes is still an active field (He et al., 2016; Yang et al., 2021). Given the constraints of having to choose specific marker genes, it is difficult for staining-based characterizations to provide tropism profiles that are relevant for diverse and changing research needs. This highlights the importance of using unbiased, full transcriptome profiling for vector characterization.

We have shown that our combined experimental and computational platform is able to recover transduction biases and profile multiple variants in a single animal, even amidst the noise of ambient RNA. We have further shown that our method is robust to the variability inherent in delivery and extraction from different animals, with different transgenes, and with different regulatory elements. For example, we discovered lower overall expression from vectors carrying UBC-mCherry compared with CAG-mNeonGreen-WPRE. Such differences are not surprising since the WPRE is known to increase RNA stability and therefore transcript abundance (Johansen et al., 2003). Furthermore, the shift in cell-type bias may come from the UBC promoter, as even ubiquitous promoters such as CAG and UBC have been shown to have variable levels of expression in different cell types (Qin et al., 2010). Despite these biases, looking at the differences in transduction between variants delivering the same construct within an individual animal reveals the strongest candidate vectors for on-target and off-target cell types of interest. While we show that our method can profile AAVs carrying standard fluorescent cargo, caution is needed when linking differences in absolute viral tropism to changes in capsid composition alone without considering the contribution of the transgene and regulatory elements. Therefore, for more robust and relative tropism between variants, we found it beneficial to use small barcodes and co-injections of pools of vectors. Our scRNA-seq-based approach is not restricted to profiling capsid variants, but can be expanded in the future to screen promoters (Chuah et al., 2014; Jüttner et al., 2019; Rincon et al., 2015), enhancers (Hrvatin et al., 2019; Mich et al., 2020), or transgenes (Gustafsson et al., 2004; Shirley et al., 2020), all of which are essential elements requiring optimization to improve gene therapy.

Finally, we have used scRNA-seq to understand how intra-orbital administration of AAV-PHP.eB affects the host cell transcriptome across distinct time points. Results from our study show genes pertaining to the p53 pathway in endothelial cells are differentially expressed 3 days after injection. The highest number of differentially expressed genes being in endothelial cells suggests that vascular cells could be the initial responders to viral transduction and expression of the transgene. This is supported by Kodali et al., who have shown that endothelial cells are the first to elicit a response to peripheral inflammatory stimulation by transcribing genes for proinflammatory mediators and cytokines (Kodali et al., 2020). With regards to p53 differentially expressed genes, Ghouzzi, et al. have also shown that the genes Phdla3, Aen, and Cdkn1a were upregulated in cells infected with ZIKA virus, signifying genotoxic stress and apoptosis induction (Ghouzzi et al., 2016). Upregulation of genes such as Bax and Cdkn1a in our data hint at an initiation of apoptosis and cell cycle arrest, respectively, in response to cellular stress induced by viral transduction (Ferreira and Nagai, 2019; Zamagni et al., 2020). The reduction in the number of differential expressed genes across all cells (Figure 5 B) at day twenty-five imparts that the initial inflammatory responses did not escalate. Downregulation of Cd209a gene in perivascular macrophages in our data further implies that the AAV-PHP.eB infection did not necessitate a primary adaptive immune response. Additionally, antigen presenting genes, such as H2-D1, returning back to control expression levels and a lack of proinflammatory cytokines being upregulated supports that the event of infiltration of peripheral leukocytes is unlikely, in agreement with prior studies (Chamberlin et al., 1998; McCown et al., 1996). Upregulation of genes such as Gadd45g, Gadd45b, and Ppp1r15a suggest that neurons are turning on stress-related programs as an early response to encountering the virus. Genes such as Nr4a1 and Dusp1, which play a role in the MAPK pathway, indicate sustained stress response even at day 25. Based on prior studies, we speculate that the genes that are differentially expressed at day 25 in the excitatory neurons are due to transgene expression and not due to the virion (Lowenstein et al., 2007). It is important to note that the findings discussed here are specific to the rAAV, transgene, and dosage. Our results highlight the power of single-cell profiling in being able to ascertain cell-type-specific responses at an early time point post-injection.

In summary, our platform could aid the gene therapy field by allowing more thorough characterization of existing and emerging recombinant AAVs by helping uncover cellular responses to rAAV-mediated gene therapy, and by guiding the engineering of novel AAV variants.

## Acknowledgements

We thank the Gradinaru and Thomson labs for helpful discussions, Allan-Hermann Pool for advice on the mouse brain tissue dissociation procedure, Jeff Park for advice on 10X Genomics Chromium single-cell library preparation, Min Jee Jang for help in designing probes and troubleshooting FISH-HCR, and Ben Deverman and Ken Chan for early discussions on strategy. This work was supported by the NIH Pioneer DP1OD025535, Beckman Institute for CLARITY, Optogenetics and Vector Engineering Research at Caltech, the Single-Cell Profiling and Engineering Center (SPEC) in the Beckman Institute at Caltech, and the Curci Foundation. V.G. and M.T. are Heritage Principal Investigators supported by the Heritage Medical Research Institute.

## Author contributions

D.B., M.A., T.D., and V.G. conceived the project and designed the experiments. S.C. and M.T. provided critical single-cell RNA sequencing expertise. T.D., M.A., and D.B. prepared the DNA constructs and produced virus. M.A. performed the injections, tissue dissociation, histology, imaging and image quantification. D.B. and T.D. performed the single-cell library preparation and prepared samples for sequencing. D.B. and M.A. built the data processing pipeline. D.B., M.A., T.D., and A.W. performed the analysis. All authors contributed to the MS as drafted by D.B., M.A., and V.G.. M.T. supervised single-cell RNA sequencing computational pipelines while V.G. supervised the overall project.

## Methods

### Animals

Animal husbandry and all experimental procedures involving animals were performed in accordance with the California Institute of Technology Institutional Animal Care and Use Committee (IACUC) guidelines and approved by the Office of Laboratory Animal Resources at the California Institute of Technology (animal protocol no. 1650). Male C57BL/6J mice (Stock No: 000664) used in this study were purchased from the Jackson Laboratory (JAX). AAV variants were injected i.v. into the retro-orbital sinus of 6–7 week old mice.

### Plasmids

In vivo vector characterization of AAV variant capsids was conducted using single-stranded (ss) rAAV genomes. pAAV:CAG-NLS-mNeonGreen, pAAV:CAG-NLS-mRuby2, pAAV:CAG-tdTomato, and pAAV:CAG-NLS-tdTomato constructs were adapted from previous publications (Chan et al., 2017; Ravindra Kumar et al., 2020). To introduce barcodes into the polyA region of CAG-NLS-mNeonGreen, we digested the plasmid with BglII and EcoRI, and performed Gibson assembly (E2611, NEB) to insert synthesized fragments with 7bp degenerate nucleotide sequences 89 bp upstream of the polyadenylation site. We then seeded bacterial colonies and selected and performed Sanger sequencing on the resulting plasmids to determine the corresponding barcode.

The UBC-mCherry-AAV-cap-in-cis plasmid was adapted from the rAAV-Cap-in-cis-lox plasmid from a previous publication (Deverman et al., 2016). We performed a restriction digest on the plasmid with BsmbI and SpeI to remove UBC-mCherry and retain the AAV9 cap gene and remaining backbone. We then circularized the digested plasmid using a gblock joint fragment to get a plasmid containing AAV2-Rep, AAV9-Cap, and the remaining backbone via T4 ligation. In order to insert UBC-mCherry with the desired orientation and location, we amplified its linear segment from the original rAAV-Cap-in-cis-lox plasmid. The linear UBC-mCherry-polyA segment and circularized AAV2-Rep,AAV9-cap plasmid were then both digested with HindIII and ligated using T4 ligation. In order to get the SV40 PolyA element in the proper orientation with respect to the inserted UBC-mCherry, we removed the original segment from the plasmid using AvrII and AccI enzymes and inserted AvrII, AccI treated SV40 gblock using T4 ligation to get the final plasmid.

To insert barcodes into UBC-mCherry-AAV-cap-in-cis, we obtained 300 bp DNA fragments containing the two desired capsid mutation regions for each variant and the variant barcode, flanked by BsrGI and XbaI cut sites. The three segments of the fragment were separated by BsaI Type I restriction sites. We digested the UBC-mCherry-AAV-cap-in-cis plasmid with BsrGI and XbaI, and ligated each variant insert to this backbone. Then, to reinsert the missing regions, we performed Golden Gate assembly with two inserts and BsaI-HF.

### Viral production

To produce viruses carrying *in trans* constructs, we followed established protocols for the production of rAAVs (Challis et al., 2019). In short, HEK293T cells were triple transfected using polyethylenimine (PEI) with three plasmids: pAAV (see Plasmids), pUCmini-iCAP-PHP.eB (Chan et al., 2017), pUCmini-iCAP-CAP-B10 (Flytzanis et al., 2020), or pUCmini-iCAP-PHP.V1 (Ravindra Kumar et al., 2020), and pHelper. After 120 h, virus was harvested and purified using an iodixanol gradient (Optiprep, Sigma). For our 7-variant pool, we modified the protocol to be a double transfection using PEI with two plasmids: UBC-mCherry-AAV-cap-in-cis and pHelper.

### Tissue processing for single-cell suspension

Three to four weeks after the injection, mice (9-10 weeks old) were briefly anesthetized with isoflurane (5%) in an isolated plexiglass chamber followed by i.p. injection of euthasol (100 mg/kg). The following dissociation procedure of cortical tissue into a single-cell suspension was adapted with modifications from a previous report (Pool et al., 2020). Animals were transcardially perfused with ice-cold carbogenated (95% O_2_ and 5% CO_2_) NMDG-HEPES-ACSF (93 mM NMDG, 2.5 mM KCl, 1.2 mM NaH_2_PO_4_, 30 mM NaHCO_3_, 20 mM HEPES, 25 mM glucose, 5 mM Na L-ascorbate, 2 mM thiourea, 3 mM Na-pyruvate, 10 mM MgSO_4_, 1 mM CaCl_2_, 1 mM kynurenic acid Na salt, pH adjusted to 7.35 with 10N HCl, osmolarity range 300–310 mOsm). Brains were rapidly extracted and cut in half along the anterior-posterior axis with a razor blade. Half of the brain was used for IHC histology while the second half of the brain was used for scRNA-seq. Tissue used for scRNA-seq was immersed in ice-cold NMDG-HEPES-ACSF saturated with carbogen. The brain was sectioned into 300-μm slices using a vibratome (VT-1200, Leica Biosystems, IL, USA). Coronal sections from Bregma −0.94 mm to −2.80 mm were collected in a dissection dish on ice containing NMDG-HEPES-ACSF. Cortical tissue from the dorsal surface of the brain to ∼3.5 mm ventral was cut out and further sliced into small tissue pieces. NMDG-HEPES-ACSF was replaced by trehalose-HEPES-ACSF (92 mM NaCl, 2.5 mM KCl, 1.2 mM NaH_2_PO_4_, 30 mM NaHCO_3_, 20 mM HEPES, 25 mM glucose, 2 mM MgSO_4_, 2 mM CaCl_2_, 1 mM kynurenic acid Na salt, 0.025 mM D-(+)-trehalose dihydrate*2H_2_O, pH adjusted to 7.35, osmolarity ranging 320–330 mOsm) containing papain (60 U/ml; P3125, Sigma Aldrich, pre-activated with 2.5 mM cysteine and a 0.5–1 h incubation at 34°C, supplemented with 0.5 mM EDTA) for the enzymatic digestion. Under gentle carbogenation, cortical tissue was incubated at 34°C for 50 min with soft agitation by pipetting every 10 min. 5 μl 2500 U/ml DNase I (04716728001 Roche, Sigma Aldrich) was added to the single-cell suspension 10 min before the end of the digestion. The solution was replaced with 200 μl trehalose-HEPES-ACSF containing 3 mg/ml ovomucoid inhibitor (OI-BSA, Worthington) and 1 μl DNase I. At room temperature, the digested cortical tissue was gently triturated with fire-polished glass Pasteur pipettes for three consecutive rounds with decreasing pipette diameters of 600, 300, and 150 μm. 800 μl of trehalose-HEPES-ACSF with 3 mg/ml ovomucoid inhibitor was added. The uniform single-cell suspension was pipetted through a 40 μm cell strainer (352340, Falcon) into a new microcentrifuge tube followed by centrifugation at 300 g for 5 min at 4°C. The supernatant was discarded and cell pellet was resuspended in 1 ml of trehalose-HEPES-ACSF. After mixing using a Pasteur pipette with a 150 μm tip diameter, the single-cell suspension was centrifuged again. Supernatant was replaced with fresh trehalose-HEPES-ACSF and the resuspended cell pellet was strained with a 20 μm nylon net filter (NY2004700, Millipore). After resuspension in trehalose-HEPES-ACSF, cells were pelleted again and resuspended in 100 μl of ice-cold resuspension-ACSF (117 mM NaCl, 2.5 mM KCl, 1.2 mM NaH_2_PO_4_, 30 mM NaHCO_3_, 20 mM HEPES, 25 mM glucose, 1 mM MgSO_4_, 2 mM CaCl_2_, 1 mM kynurenic acid Na salt and 0.05% BSA, pH adjusted to 7.35 with Tris base, osmolarity range 320–330 mOsm). Cells were counted with a hemocytometer and the final cell densities were verified to be in the range of 400–2,500 cells/μl. The density of single-cell suspension was adjusted with resuspension-ACSF if necessary.

### Transcriptomic library construction

Cell suspension volumes containing 16,000 cells–expected to retrieve an estimated 10,000 single-cell transcriptomes–were added to the 10x Genomics RT reaction mix and loaded to the 10x Single Cell Chip A (230027, 10x Genomics) for 10x v2 chemistry or B (2000168, 10x Genomics) for 10x v3 chemistry per the manufacturer’s protocol (Document CG00052, Revision F, Document CG000183, Revision C, respectively). We used the Chromium Single Cell 3’ GEM and Library Kit v2 (120237, 10x genomics) or v3 (1000075, 10x Genomics) to recover and amplify cDNA, applying 11 rounds of amplification. We took 70 ng to prepare Illumina sequencing libraries downstream of reverse transcription following the manufacturer’s protocol, applying 13 rounds of sequencing library amplification.

### Viral library construction

We selectively amplified viral transcripts from 15 ng of cDNA using a cargo-specific primer binding to the target of interest and a primer binding the partial Illumina Read 1 sequence present on the 10x capture oligos (Supplemental Table 1). For animals injected with a single cargo, amplification was performed only once using the primer for the delivered cargo; for animals with distinct cargo sequences per variant, amplification was performed in parallel reactions from the same cDNA library using different cargo-specific primers for each reaction. We performed the amplification using 2x KAPA HiFi HotStart ReadyMix (KK2600) for 28 cycles at an annealing temperature of 53°C. Afterwards, we performed a left-sided SPRI cleanup with a concentration dependent on the target amplicon length, in accordance with the manufacturer’s protocol (SPRISelect, Beckman Coulter B23318). We then performed an overhang PCR on 100 ng of product with 15 cycles using primers that bind the cargo and the partial Illumina Read 1 sequence and appending the P5/P7 sequences and Illumina sample indices. We performed another SPRI cleanup, and analyzed the results via an Agilent High Sensitivity DNA Chip (Agilent 5067-4626).

### Sequencing

Transcriptome libraries were pooled together in equal molar ratios according to their DNA mass concentration and their mean transcript size as determined via bioanalyzer. Sequencing libraries were processed on Novaseq 6000 S4 300-cycle lanes. The run was configured to read 150 bp from each end. Sequencing was outsourced to Fulgent Genetics and the UCSF Center for Advanced Technology.

All viral transcript libraries except barcoded UBC-mCherry were pooled together in equal molar ratios into a 4 nM sequencing library, then diluted and denatured into a 12 pM library as per the manufacturer’s protocol (Illumina Document #15039740v10). The resulting library was sequenced using a MiSeq v3 150-cycle reagent kit (MS-102-3001), configured to read 91 base pairs for Read 2 and 28 base pairs for Read 1. To characterize the effect of sequencing depth, one viral transcript library was additionally processed independently on a separate MiSeq run.

The UBC-mCherry viral transcript library, which was recovered with primers near the polyadenylation site, consisted of fragments ∼307 bp long. Since this length is within the common range for an Illumina NovaSeq run, this viral transcript library was pooled and included with the corresponding transcriptome library.

### Transcriptome read alignment

For transcriptome read alignment and gene expression quantification, we used 10x Cell Ranger v5.0.1 with default options to process the FASTQ files from the transcriptome sequencing library. The reads were aligned against the mus musculus reference provided by Cell Ranger (mm10 v2020-A, based on Ensembl release 98).

To detect viral transcripts in the transcriptome, we ran an additional alignment using 10x Cell Ranger v5.0. 1 with a custom reference genome based on mm10 v2020-A. We followed the protocol for constructing a custom Cell Ranger reference as provided by 10x Genomics. This custom reference adds a single gene containing all the unique sequences from our delivered plasmids in the study, delineated as separate exons. Sequences that are common between different cargo are provided only once, and annotated as alternative splicings.

### Viral transcript read alignment

For viral read alignment, we aligned each Read 2 to a template derived from the plasmid, excluding barcodes. The template sequence was determined by starting at the ATG start site of the XFP cargo and ending at the AATAAA polyadenylation stop site. We used a Python implementation of the Striped Smith-Waterman algorithm from scikit-bio to calculate an alignment score for each read, and normalized the score by dividing by the maximum possible alignment score for a sequence of that length, minus the length of the barcode region. For each Read 2 that had a normalized alignment score of greater than 0.7, we extracted the corresponding cell barcode and UMI from Read 1, and any insertions into the template from Read 2.

### Constructing the variant lookup table

For co-injections with multiple templates and injections of barcoded templates, we constructed a lookup table to identify which variant belongs to each cell barcode/UMI. For each template, we counted the number of reads for each cell barcode/UMI. For reads of barcoded cargo, we only counted reads where the detected insertion in the barcode region unambiguously aligned to one of the pre-defined variant barcodes. Due to sequencing and PCR amplification errors, most cell barcode/UMI combinations had reads associated with multiple variants. Thus, we identified the variant with the largest count for each cell barcode/UMI. We discarded any cell barcode/UMIs that had more than one variant tied for the largest count. Finally, each cell barcode/UMI that was classified as a viral transcript in the transcriptome (see Transcriptome read alignment) was converted into the virus detected in the variant lookup table, or was discarded if it did not exist in the variant lookup table.

### Estimating transduction rate

To determine an estimate of the percent of cells within a group expressing viral cargo above background, we compared the viral transcript counts in that group of cells to a background distribution of viral transcript counts in debris (see Droplet type classification). First, we obtained the empirical distribution of viral transcript counts by extracting the viral counts for that variant in cell barcodes classified as the target cell type as well as cell barcodes classified as debris. Next, we assumed a percentage of cells containing debris. For each viral transcript count, starting at 0, we calculated the number of cells that would contain this transcript count, if the assumed debris percentage was correct. We then calculated an error between this estimate and the number of cells with this transcript count in the cell type of interest. We tallied this error over all the integer bins in the histogram, allowing the error in a previous bin to roll over to the next bin. We repeated this for all possible values of percentage of debris from 0 to 100 in increments of 0.25, and the value that minimized the error was the estimated percentage of cells whose viral transcript count could be accounted for by debris. The inverse of this was our estimate of the number of cells expressing viral transcripts above background.

To validate that this method reliably recovers an estimate of transduction rate, we performed a series of simulations using models of debris viral transcript counts added to proposed cell type transcript count distributions across a range of parameterizations. To get estimates of the background distribution of debris, we used diffxpy (https://github.com/theislab/diffxpy) to fit the parameters of a negative binomial distribution to the viral transcript counts in debris droplets within a sample. We then postulated 1,000 different parameterizations of the negative binomial representing transcript counts in groups of cells, with 40 values of r ranging from 0.1 to 10, spaced evenly apart, and 25 values of p ranging from 0.001 to 0.99, spaced evenly apart. For each proposed negative binomial model, we drew 1,000 random samples of viral counts from the learned background distribution, and 1,000 random samples from the proposed cell distribution, and summed the two vectors. This summed vector was then used in our transduction rate estimation function, along with a separate 1,000 random samples of background viral transcripts for the function to use as an estimate of the background signal. We calculated the true probability of non-zero expression in our proposed cell negative binomial model (1 – P(X = 0)), and compared this value with the estimated value from the transduction rate estimation method.

### Calculating viral tropism

For each variant *v_n_* and cell type of interest *c_i_*, we estimated the percentage of cells expressing viral cargo. To calculate tropism bias, we used this estimated expression rate, 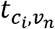, to estimate the number of cells expressing viral transcripts in that cell type, 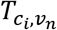 out of the total number of cells of that type, 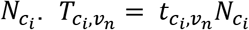. Cell type bias, 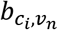 within a sample was then calculated as the ratio of the number of cells of interest divided by the total number of transduced cells, 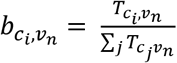. Finally, to calculate the difference in transduction bias for a particular variant relative to other variants in the sample, 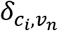, we subtracted the bias of the variant from the mean bias across all other variants, 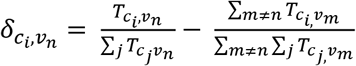.

### Histology

#### Immunohistochemistry

The immunohistochemistry procedure was adapted from a previous publication (Oikonomou et al., 2019). Brain tissue was fixed in 4% paraformaldehyde (PFA) at 4°C overnight on a shaker. Samples were immersed in 30% sucrose in 1x phosphate buffered saline (PBS) solution for >2 days and then embedded in Tissue-Tek O.C.T. Compound (102094-104, VWR) before freezing in dry ice for 1 h. Samples were sectioned into 50 μm coronal slices on a cryostat (Leica Biosystems). Brain slices were washed once with 1x phosphate buffered saline (PBS) to remove O.C.T. Compound. Samples were then incubated overnight at 4°C on a shaker in a 1x PBS solution containing 0.1% Triton X-100, 10% normal goat serum (NGS; Jackson ImmunoResearch, PA, USA), and primary antibodies. Sections were washed three times for 15 min each in 1x PBS. Next, brain slices were incubated at 4°C overnight on a shaker in a 1x PBS solution containing 0.1% Triton X-100, 10% NGS, and secondary antibodies. Sections were washed again three times for 15 min each in 1x PBS. Finally, slices were mounted on glass microscope slides (Adhesion Superfrost Plus Glass Slides, #5075-Plus, Brain Research Laboratories, MA, USA). After the brain slices dried, DAPI-containing mounting media (Fluoromount G with DAPI, 00-4959-52, eBioscience, CA, USA) was added before protecting the slices with a cover glass (Cover glass, #4860-1, Brain Research Laboratories, MA, USA). Confocal images were acquired on a Zeiss LSM 880 confocal microscope (Zeiss, Oberkochen, Germany). The following primary antibodies were used: rabbit monoclonal to NeuN (Rbfox3) (1:500; ab177487; Abcam, MA, USA), rabbit monoclonal to S100 beta (1:500; ab52642; Abcam, MA, USA), and rabbit monoclonal to Olig2 (1:500; ab109186; Abcam, MA, USA). The following secondary antibody was used: goat anti-rabbit IgG H&L Alexa Fluor 647 (1:500; ab150079; Abcam, MA, USA).

#### Fluorescent in situ hybridization chain reaction

FISH-HCR was conducted as previously reported (Patriarchi et al., 2018). Probes targeting neuronal markers were designed using custom-written software (https://github.com/GradinaruLab/HCRprobe). Probes contained a target sequence of 20 nucleotides, a spacer of 2 nucleotides, and an initiator sequence of 18 nucleotides. Criteria for the target sequences were: (1) a GC content between 45%–60%, (2) no nucleotide repeats more than three times, (3) no more than 20 hits when blasted, and (4) the ΔG had to be above –9 kcal/mol to avoid self-dimers. Last, the full probe sequence was blasted and the Smith-Waterman alignment score was calculated between all possible pairs to prevent the formation of cross-dimers. In total, we designed 26 probes for *Gad1*, 20 probes for *Vip*, 22 probes for *Pvalb*, 18 probes for *Sst,* and 28 probes for *Slc17a7*. Probes were synthesized by Integrated DNA Technologies.

### Droplet type identification

scRNA-seq datasets were analyzed with custom-written scripts in Python 3.7.4 using a custom fork off of scVI v0.8.1, and scanpy v1.6.0. To generate a training dataset for classifying a droplet as debris, multiplets, neuronal, or non-neuronal cells, we randomly sampled cells from all 27 cortical tissue samples. We sampled a total of 200,000 cells, taking cells from each tissue sample proportional to the expected number of cells loaded into the single-cell sequencing reaction. Within each sample, cells were drawn randomly, without replacement, weighted proportionally by their total number of detected UMIs. For each sample, we determined a lower bound on the cutoff between cells and empty droplets by constructing a histogram of UMI counts per cell from the raw, unfiltered gene count matrix. We then found the most prominent trough preceding the first prominent peak, as implemented by the scipy peak_prominences function. We only sampled from cells above this lower bound. Using these sampled cells, we trained a generative neural network model via scVI with the following parameters: 20 latent features, 2 layers, and 256 hidden units. These parameters were chosen from a coarse hyperparameter optimization centered around the scVI default values (Supplemental Table 3). We included the sample identifier as the batch key so that the model learned a latent representation with batch correction.

After training, Leiden clustering was performed on the learned latent space as implemented by scanpy. We used default parameters except for the resolution, which we increased to 2 to ensure isolation of small clusters of cell multiplets. Using the learned generative model, we draw 5000 cells from the posterior distribution based on random seed cells in each cluster. We draw an equal number conditioned on each batch. From these samples, we then calculated a batch-corrected probability of each cluster expressing a given marker gene (see Cluster marker gene determination). For this coarse cell typing, we chose a single marker gene for major cell types expected in the cortex (Supplemental Table 2). If a cluster was expressing the neuron marker gene Rbfox3, it was labeled as “Neurons”. If a cluster was expressing any of the other non-neuronal marker genes, it was labelled as “Non-neurons”. Next, we ran Scrublet on the training cells to identify potential multiplets. Scrublet was run on each sample independently, since it is not designed to operate on combined datasets with potential batch-specific confounds. We then calculated the percentage of droplets in each cluster of the combined data that were identified as multiplets by Scrublet. We found a percentage threshold for identifying a cluster as containing predominantly multiplets by using Otsu’s threshold, as implemented by scikit-image. All droplets in any cluster above the multiplet percentage threshold were labelled as “Multiplets”. All other clusters were labelled as “Debris”.

Next, we trained a cell-type classifier using scANVI on the droplets labeled as training data. We used the weights from the previously trained scVI model as the starting weights for scANVI. Rather than using all cells for every epoch of the trainer, we implemented an alternative sampling scheme that presented each cell type to the classifier in equal proportions. Once the model was trained, all cells above the UMI lower noise bound were run through the classifier to obtain their cell-type classification. Droplets classified as “Neurons” or “Non-neurons” were additionally filtered by their scANVI-assigned probability. We retained only cells above an FDR threshold of 0.05, corrected for multiple comparisons using the Benjamini-Hochberg procedure. Finally, since the original run of Scrublet for multiplet detection was performed on only the training data, and thus did not take advantage of all the cells available, we ran Scrublet on all droplets classified as cells, and removed any identified multiplets.

### Cluster marker gene determination

To identify which clusters are expressing marker genes, we determined an estimated probability of a marker gene being expressed by a random cell in that cluster. For each cluster, we randomly sampled 5,000 cells, with replacement. We used scVI to project each cell into its learned latent space, and then used scVI’s posterior predictive sampling function to generate an example cell from this latent representation, and tallied how many times the gene is expressed. We repeated this for each batch, conditioning the posterior sample on that batch, to account for technical artifacts such as sequencing depth. Once we obtained a probability of expression of a marker gene for each cluster, we find a threshold for expression using Otsu’s method, as implemented by scikit-image. Clusters that have a probability of expression above the threshold are considered positive for that marker gene.

### Neuronal subtype classification

Cells classified as neurons were further subtyped using annotations from a well-curated reference dataset. We used the Mouse Whole Cortex and Hippocampus 10x dataset from the Allen Institute for Brain Science as our reference dataset (Yao et al., 2021). First, we filtered the reference dataset to contain only cell types that are found within the brain regions collected for our experiments. To ensure that, overall, enough cells per cell type were present in our datasets, we merged cell types with common characteristics, such as expression of key marker genes. We re-aligned our cell transcriptome reads to the same pre-mRNA reference used to construct the reference dataset, so that the gene count matrices had a 1:1 mapping. We then trained a joint scANVI model with all cells identified as neurons from our samples and the reference database to learn a common latent space between them. The model was trained to classify cells based on the labels provided in the reference dataset. Cells were sampled from each class in equal proportions during training. After the model was trained, all neurons from our sample were run through the model to obtain their cell type classification.

### Non-neuronal subtype classification

Cells classified as non-neuronal were further subtyped using automatic clustering and marker gene identification. We trained an scVI model using only the non-neuronal cells and performed Leiden clustering as implemented by scanpy on the latent space. We determined which clusters were expressing each of 31 marker genes across 13 cell subtypes. Marker genes were identified from a review of existing scRNA-seq, bulk RNA-seq, or IHC studies of mouse brain non-neuronal subtypes (Supplemental Table 2). Each cluster was assigned to a cell subtype if it was determined positive for all the marker genes for that cell subtype (see Cluster marker gene determination). If a cluster contained all the marker genes for multiple cell subtypes, the cluster was assigned to the cell subtype with the greatest number of marker genes. Clusters that did not express all the marker genes for any cell subtype were labeled as “Unknown”. Clusters that expressed all the marker genes for multiple cell subtypes with the same total number of marker genes were labeled as “Multiplets”. For cell types that contained multiple clusters, we then calculated the probability of every gene being zero in each cluster (see Cluster marker gene determination). We then compared gene presence between clusters of the same cell type to see if there were any subclusters that had a dominant marker gene (present in > 50% of samples), that was not present in any of the other clusters (< 10% of samples). For the three cell types that had unique marker genes, we named the cluster after the gene with the highest 2-proportion z-score between the sampled gene counts in that cluster vs the rest.

### Quantification of images

Quantitative data analysis of confocal images was performed blind with regard to AAV capsid variant. Manual quantification was performed using the Cell Counter plugin, present in the Fiji distribution of ImageJ (National Institutes of Health, Bethesda, MD) (Schindelin et al., 2012). Transduction rate was calculated as the total number of double positive cells (i.e. viral transgene and cell type marker) divided by the total number of cell type marker labeled cells. For each brain slice, at least 100 cells positive for the gene markers of interest were counted in the cortex.

### Differential expression

To calculate differential expression within cell types between groups of animals, we used the DESeq2 R package (Love et al., 2014). For each cell type, the gene counts are summed across all cells of that type and treated as a pseudo-bulk sample. The summed gene counts from each animal are then included as individual columns for a DESeq2 differential expression analysis. We performed 3 DPI DE and 25 DPI separately, testing each sample against saline-injected controls. For each cell type, only genes that were present in all samples of at least one condition are included.

### Marker gene dot plots

To generate dot plots for marker genes, we used scanpy’s dotplot function (Wolf et al., 2018). Gene counts were normalized to the sum of the total transcript counts per cell using scanpy’s normalize_total function. Normalized gene expression values are log-transformed as part of the plotting function.

### Statistics

Statistical analyses comparing the fraction of transduced cells in different cell types for Figures 2, 3, and 4 C were conducted using GraphPad Prism 9. Statistical analyses comparing proportions of transduced cells within an animal in Figure 4 E and Figure 6 were performed using the Python statsmodels library v0.12.1. No statistical methods were used to predetermine sample sizes. The statistical test applied, sample sizes, and statistical significant effects are reported in each figure legend. The significance threshold was defined as a = 0.05.

## Supplemental Figures

**Supplemental Figure 1.**
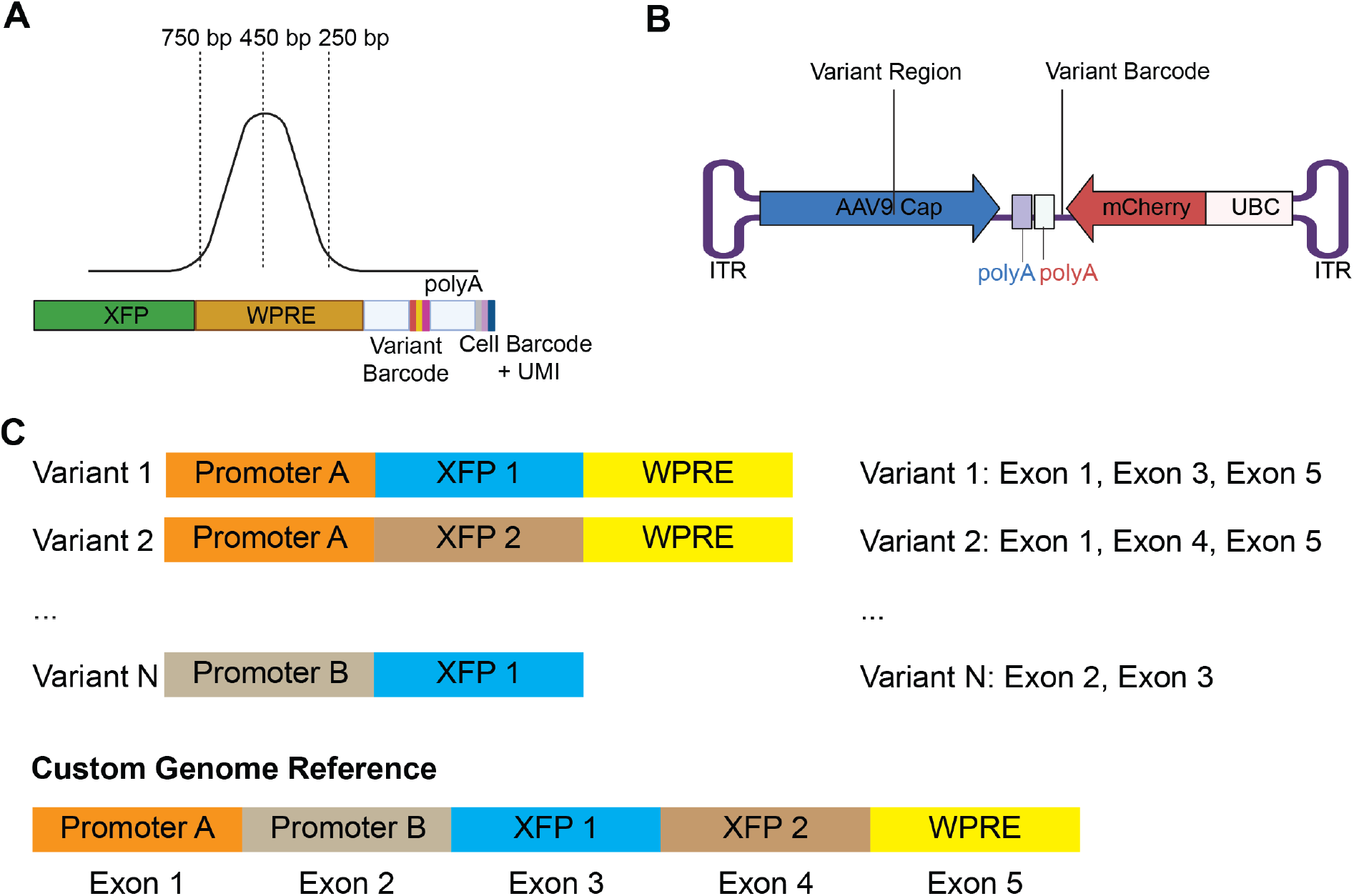
Plasmid details. (A) Size of typical transcriptome cDNA library post-fragmentation. Both distinguishing XFPs and variant barcodes fall outside the typical capture region of single-cell RNA sequencing workflows. (B) UBC-mCherry-AAV-cap-in-cis plasmid used for 7-variant barcoded pool. (C) Visualization of the construction procedure for the custom genome reference. Variant cargos are segmented into common and uncommon regions, and each unique segment is concatenated together as a contiguous gene. Variants are defined as different splicings of the custom AAV gene.

**Supplemental Figure 2.**
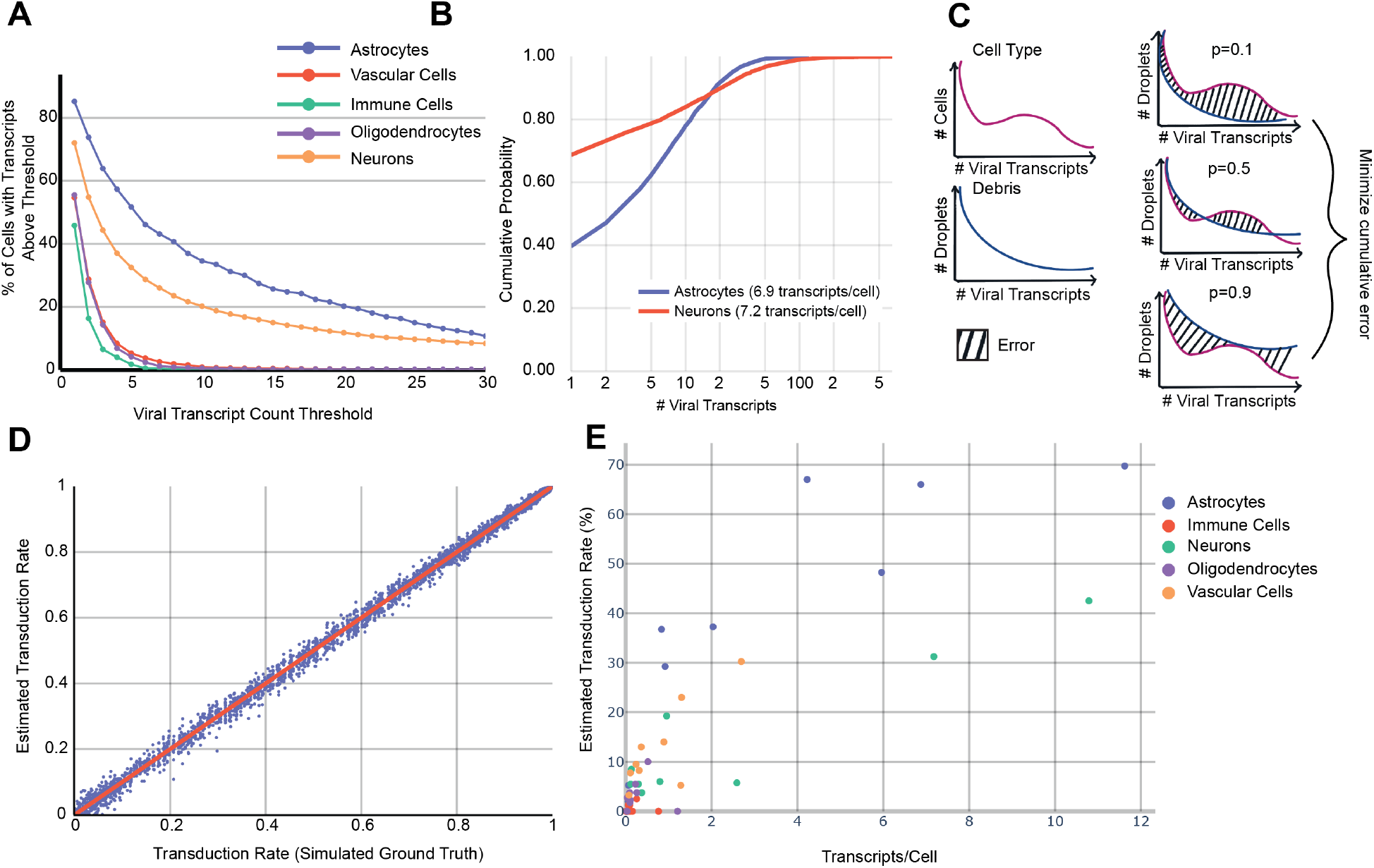
Expression rate estimation. (A) Percent of cells expressing AAV-PHP.eB cargo transcripts above a fixed threshold in a single sample. (B) An example of the distribution of viral transcript counts in a single animal from AAV-PHP.eB carrying CAG-mNeonGreen-WPRE in neurons and astrocytes. (C) Visualization of our expression-rate estimation algorithm. The distribution of the cell type of interest and background debris is obtained. An error is calculated for different estimates of the percent of the cells that express background levels of transcripts. This error is minimized to find the best fit. (D) Performance of the expression rate estimation algorithm on simulated data consisting of negative binomial distributions with parameters r between 0.1 and 10 and p between 0.001 and 0. 99, spaced evenly apart. (E) Comparison between mean transcripts/cell (x) and the estimated transduction rate (y) in major cell types for AAV-PHP.eB across 9 samples.

**Supplemental Figure 3.**
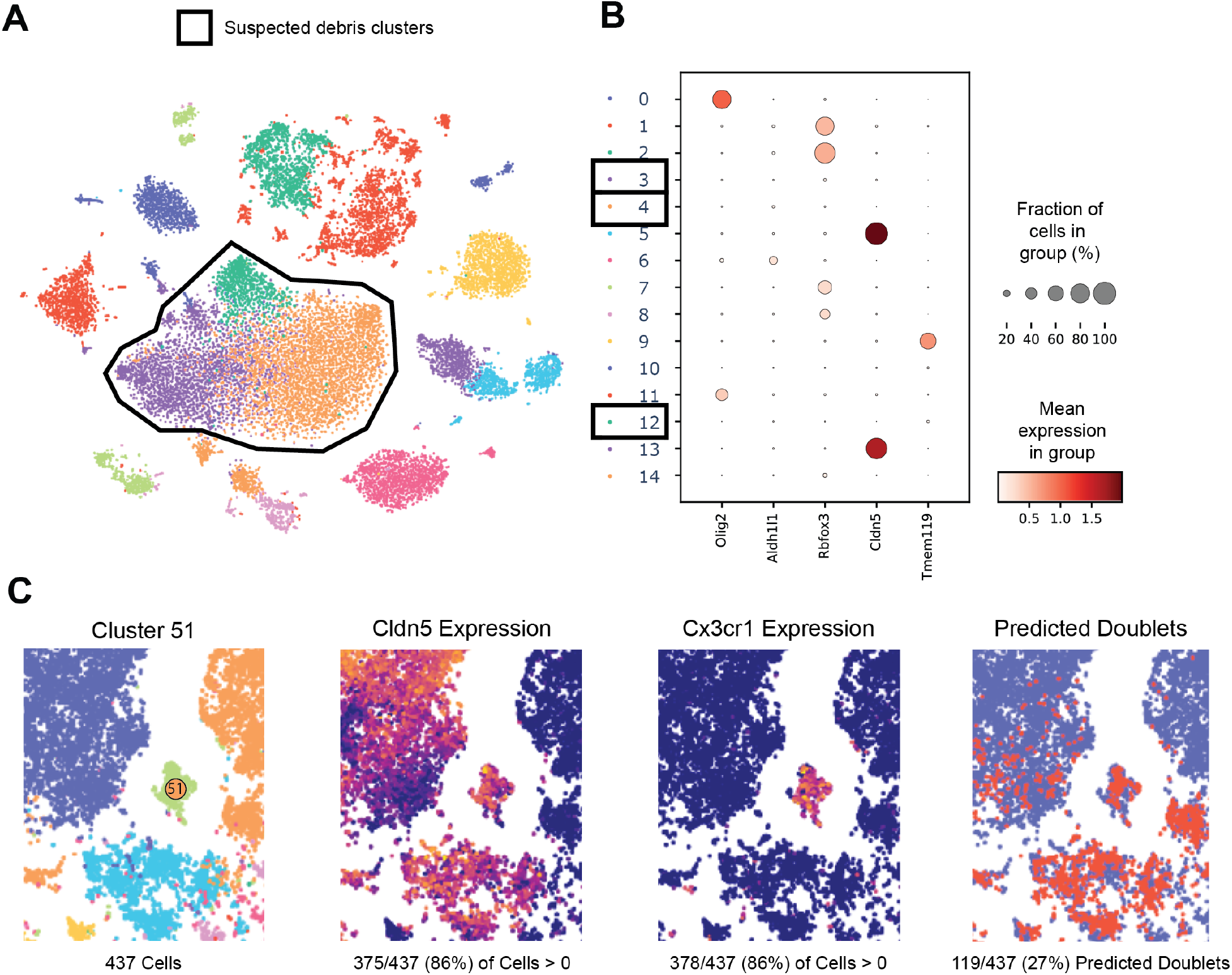
Noise from debris and doublets. (A) An example of a Cell Ranger filtered dataset. This is a t-SNE projection of the log-normalized gene expression space. Suspected debris clusters are outlined. (B) Marker gene expression for the major cell types in the brain—Oligodendrocytes/Olig2, Astrocytes/Aldh1l1, Neurons/Rbfox3, Vascular Cells/Cldn5, Immune Cells/Tmem119—for each cluster. Darker colors indicate higher mean expression, and dot size correlates with the abundance of the gene in that cluster. (C) An example of a multiplet cluster from the joint scVI space of all training samples, projected via t-SNE. Cluster 51 is annotated, and raw gene expression of Cldn5 and Cx3cr1 are shown. The percentage of cells in cluster 51 expressing each marker gene is displayed. (right) Predicted doublets from Scrublet are overlaid in red.

**Supplemental Figure 4.**
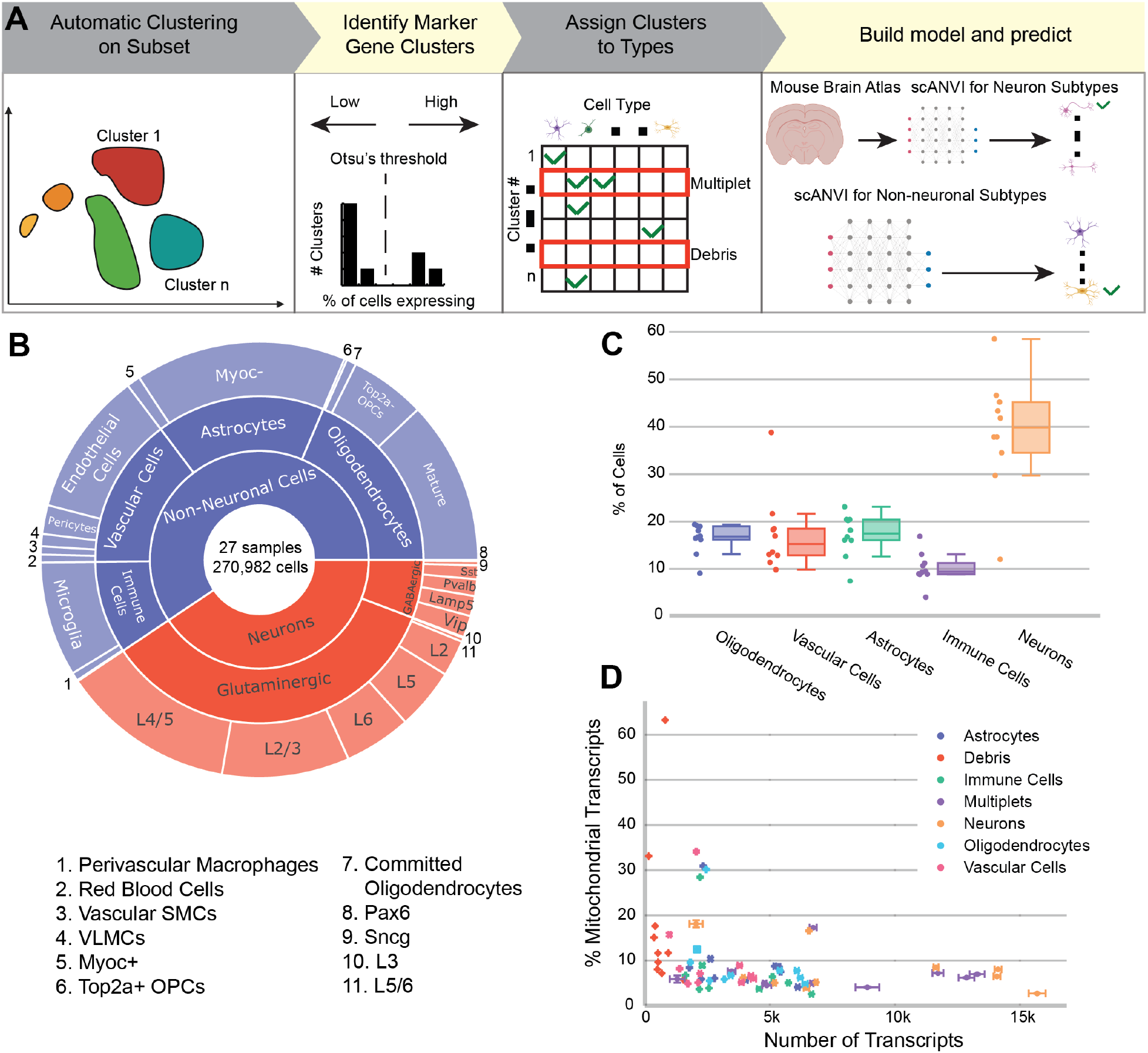
Cell typing. (A) Cell typing workflow. A subset of cells are used for training. For each marker gene, clusters expressing that marker gene are identified. Clusters that have no marker genes (debris) or are determined to be multiplets via Scrublet are marked for removal. Training data used to train a scANVI model to predict the remaining cells. A reference database can be used instead of manually labeled cells, as we did for neuronal subtypes. (B) Cell-type distribution of all identified cells from our combined cell-type taxonomy. This includes samples described in the study as well as additional controls and animals used for troubleshooting and prototyping. (C) Cell-type percentages across the major cell types in the ten samples used for AAV tropism characterization. One of the samples, BC1, had dramatically fewer neurons than any other sample and correspondingly higher percentages of non-neurons. (D) Mitochondrial gene ratio and total transcript counts of the major cell type clusters in the ten samples used for tropism characterization.

**Supplemental Figure 5.**
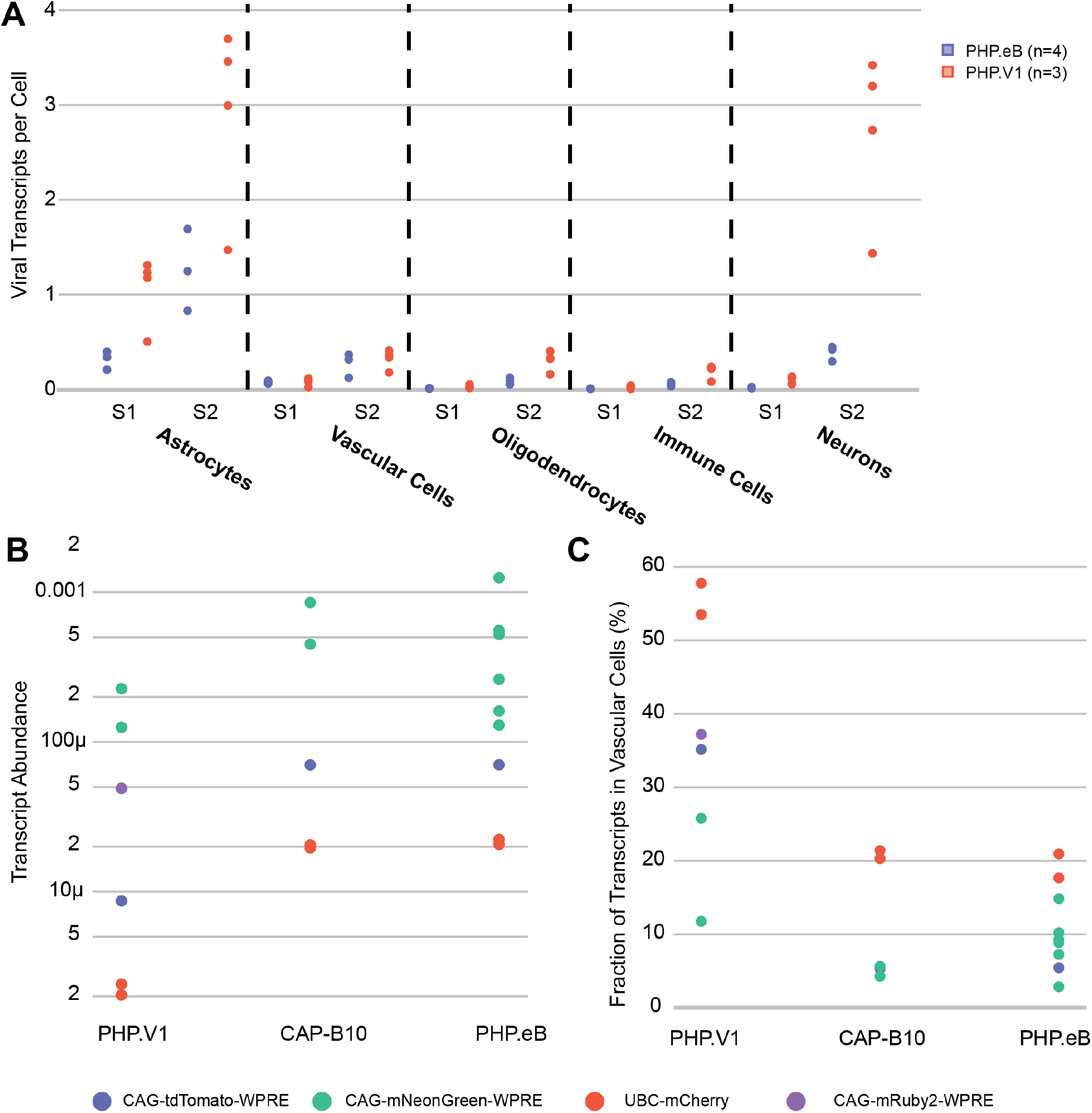
Transcript expression. (A) Viral transcript expression of different barcodes across two samples (S1, S2). Each point is a distinct barcode. (B) Viral transcript abundance in entire samples (viral transcripts / total transcripts) across different variants carrying different cargo. (C) Fraction of transcripts detected in vascular cells vs all other cell types.

**Supplemental Figure 6.**
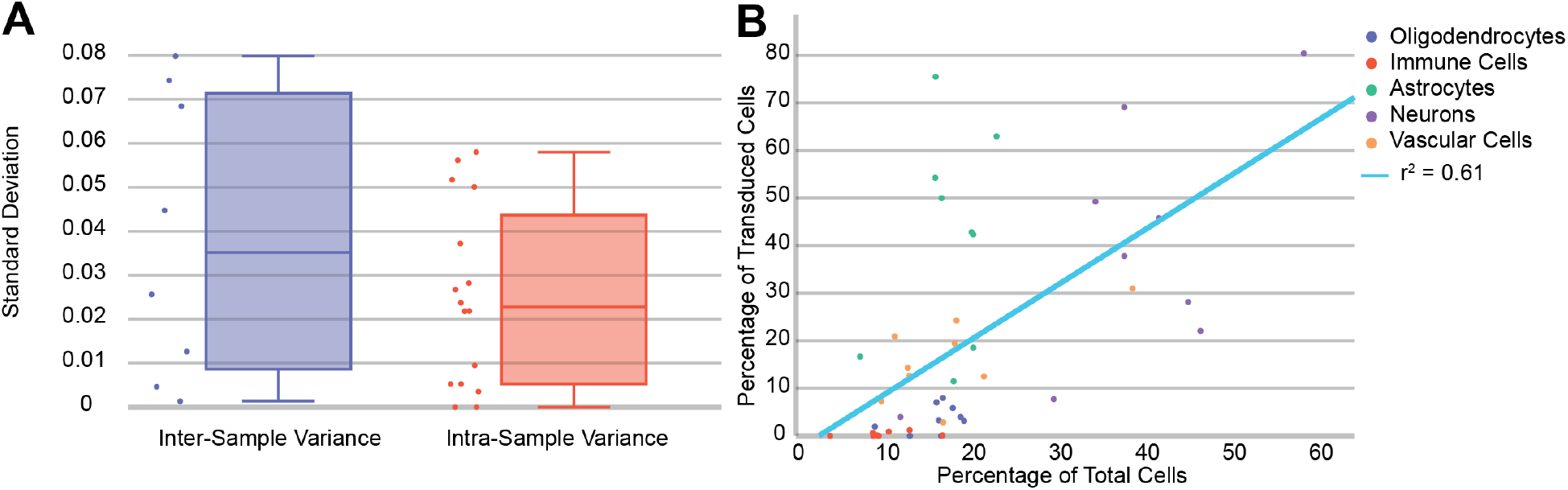
Inter-sample variability. (A) The standard deviations between measurements of the fraction of transduced cells in all major non-neuronal cell types in AAV-PHP.V1 and AAV-PHP.eB. Inter-sample variance (left) refers to the standard deviation between animals, and intra-sample variance (right) refers to the standard deviation between barcodes within the same animal. (B) The distribution of recovered cell types compared to the distribution of transduced cells across nine samples injected with AAV-PHP.eB.

**Supplemental Figure 7.**
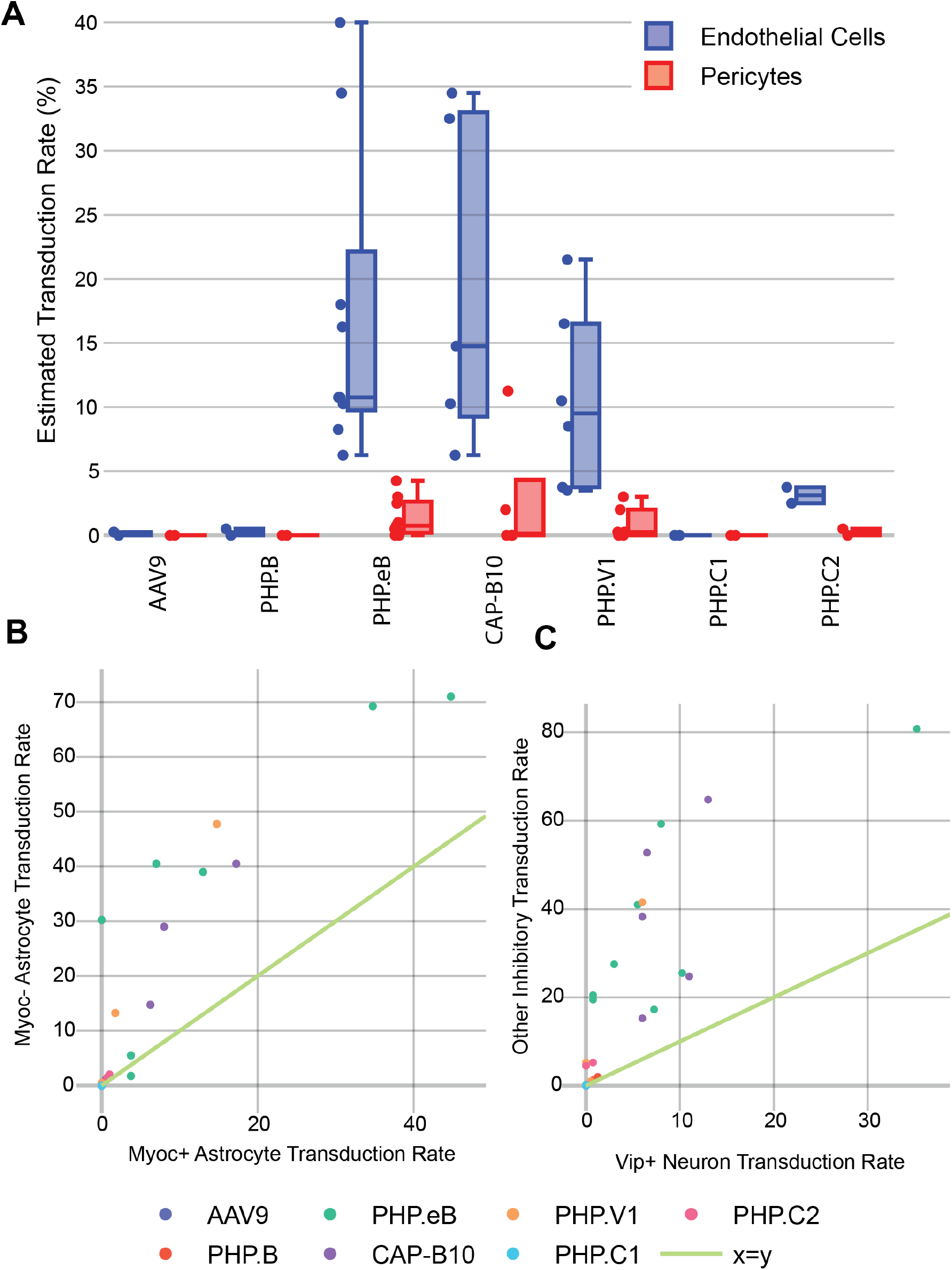
Cell subtype inspection. (A) Estimated transduction rate of endothelial cells vs pericytes across all samples and variants. (B) Pairwise transduction rate of Myoc+ and Myoc-astrocytes across all variants and samples. Each point is a single variant in a different sample. (C) Pairwise transduction rate of Vip+ neurons vs all other inhibitory neurons across all variants and samples.

**Supplemental Figure 8.**
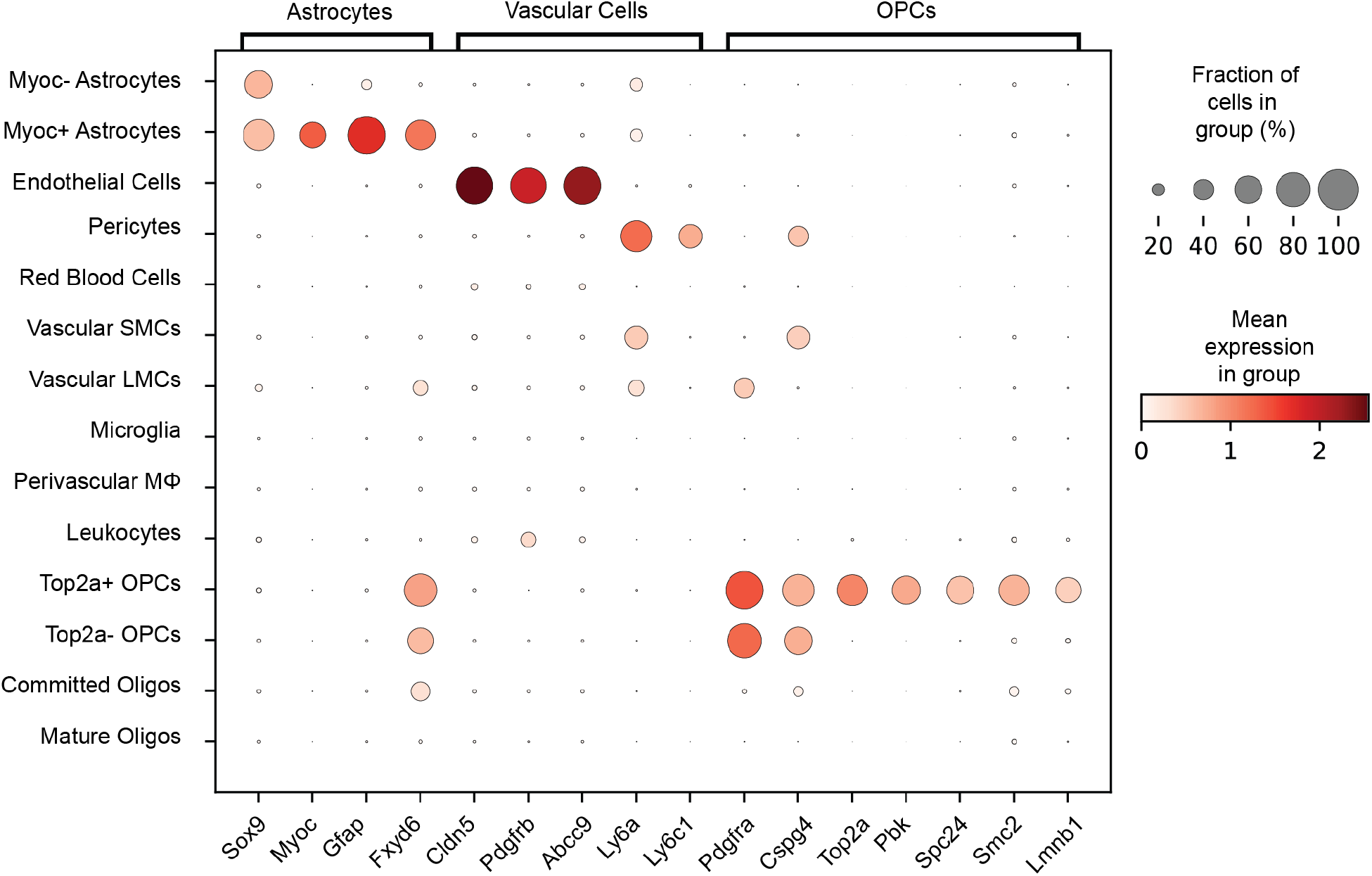
Cell subtype markers. Gene expression of additional marker genes for astrocyte and OPC subtypes.

**Supplemental Table 1.**
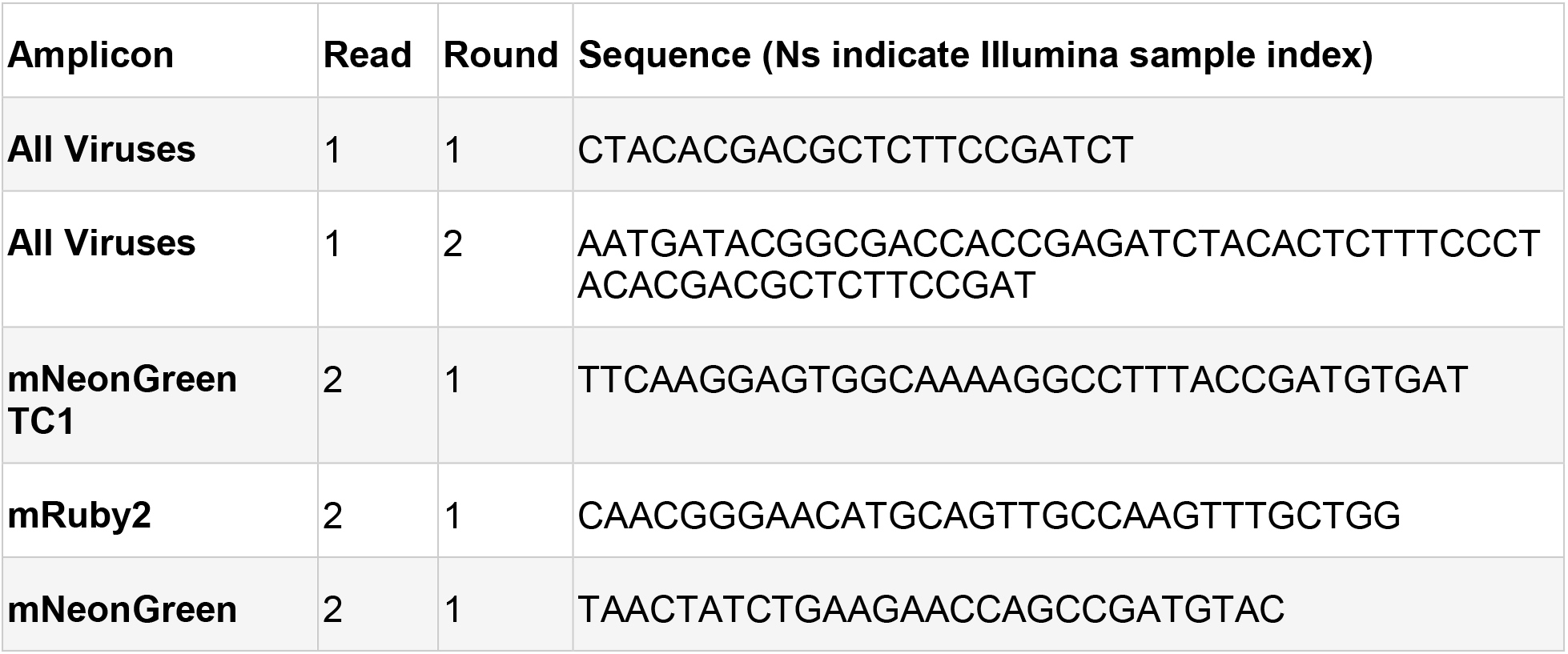
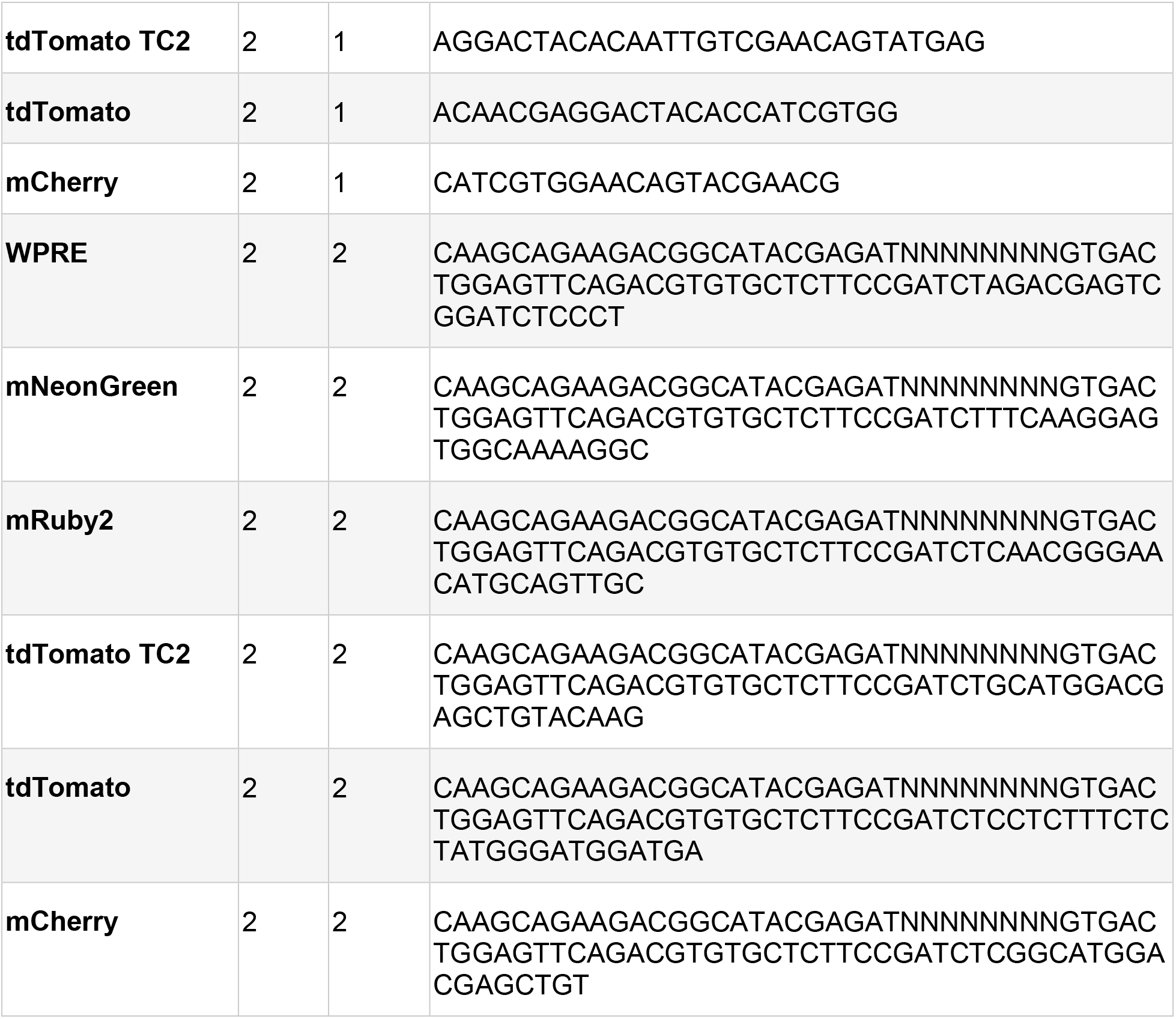
Primers. Primers used for round 1 and round 2 amplification of viral transcripts. Primers with TC1 and TC2 in the amplicon name indicate they were used only for those samples.

**Supplemental Table 2.** Marker Genes.

| **Cell Type** | **Marker Gene(s)** |
| --- | --- |
| **Astrocytes** | Aldh1l1 (*Cahoy et al., 2008*), Sox9 (*Sun et al., 2017*) |
| **Neurons** | Rbfox3 (*Lin et al., 2016*) |
| **Vascular Cells** | Cldn5 (*Song et al., 2020*) |
| **Endothelial Cells** | Slc2a1 (*Veys et al., 2020*) |
| **Pericytes** | Pdgfrb (*Winkler et al., 2010*), Rgs5, Abcc9 (*He et al., 2016*) |
| **Red Blood Cells** | Hba-a1, Hba-a2 (*Capellera-Garcia et al., 2016*) |
| **Vascular SMCs** | Acta2, Myh11, Tagln (*Chasseigneaux et al., 2018*) |
| **Vascular LMCs** | Fam180a, Slc6a13, Dcn, Ptgds (*Marques et al., 2016*) |
| **Microglia** | Cx3cr1, Tmem119 (*Jordão et al., 2019*) |
| **Leukocytes** | Itgal, Gzma (*Huang and Sabatini, 2020*) |
| **Perivascular Macropages** | Mrc1 (*Jordão et al., 2019*) |
| **Oligodendrocytes** | Olig2 (*Dai et al., 2015*) |
| **OPCs** | Pdgfra, Cspg4 (*Suzuki et al., 2017*) |
| **Mature Oligos** | Mog, Mbp (*Miron et al., 2011*) |
| **Committed Oligos** | Ptprz1, Bmp4, Nkx2-2, Vcan (*Marques et al., 2016*) |

**Supplemental Table 3.** scVI Hyperparameter Tuning.

| **Dispersion** | **Latent Lib Size** | **# Latent** | **# Layers** | **# Hidden** | **Test KL Divergence** |
| --- | --- | --- | --- | --- | --- |
| **Gene** | False | 10 | 1 | 128 | 5366.4 |
| **Gene-batch** | True | 10 | 1 | 128 | 5406.1 |
| **Gene** | True | 10 | 1 | 128 | 5391.0 |
| **Gene-batch** | True | 50 | 2 | 512 | 5362.6 |
| **Gene-batch** | False | 10 | 1 | 128 | 5378.7 |
| **Gene** | False | 25 | 1 | 128 | 5354.6 |
| **Gene** | False | 25 | 2 | 256 | 5337.8 |
| **Gene** | False | 20 | 2 | 256 | 5336.7 |
| **Gene** | False | 40 | 4 | 1024 | 5338.0 |

**Supplemental Table 4.** Sample Metadata. Supplemental file contains the following fields.

| **Field Name** | **Description** |
| --- | --- |
| **10X Version** | Whether the sample was processed using 10X V2 or V3 chemistry |
| **Animal ID** | A unique animal identifier. Some animals provided multiple samples |
| **Target # Cells** | The target number of cells for extraction. 1.6X this number is loaded into the 10X Chromium instrument |
| **# Recovered Cells** | The number of cells recovered, after debris and multiplet filtering |
| **Cell Ranger # Cells** | The number of cells as predicted by Cell Ranger |
| **Predicted Multiplets** | The number of predicted multiplets |
| **Transcriptome Sequencing Depth** | The number of reads |
| **Transcriptome Reads/Cell** | The number of reads divided by the number of recovered cells |
| **Median UMIs/Cell** | Of the recovered cells, the median total UMI count |
| **Median Genes/Cell** | Of the recovered cells, the median number of genes detected with at least one transcript |
| **Variants Recovered** | Which variants were recovered from this sample. Samples labeled “Cell Typing Only” were not used for tropism analysis, but were included in the cell type classifier |
| **Virus Sequencing Depth** | The number of reads of the amplified viral transcripts across all templates |
| **Virus Reads/Cell** | The read depth of the amplified viral transcripts |
| **Age at Extraction (Days)** | The age of the animal at extraction time |
| **Virus Incubation Time (Days)** | How many days prior to extraction the animal was injected |
| **Percent of Virus UMIs Determined** | What percent of transcriptome reads that aligned to the virus gene were disambiguated from the amplified lookup table |

**Supplemental Table 5.**
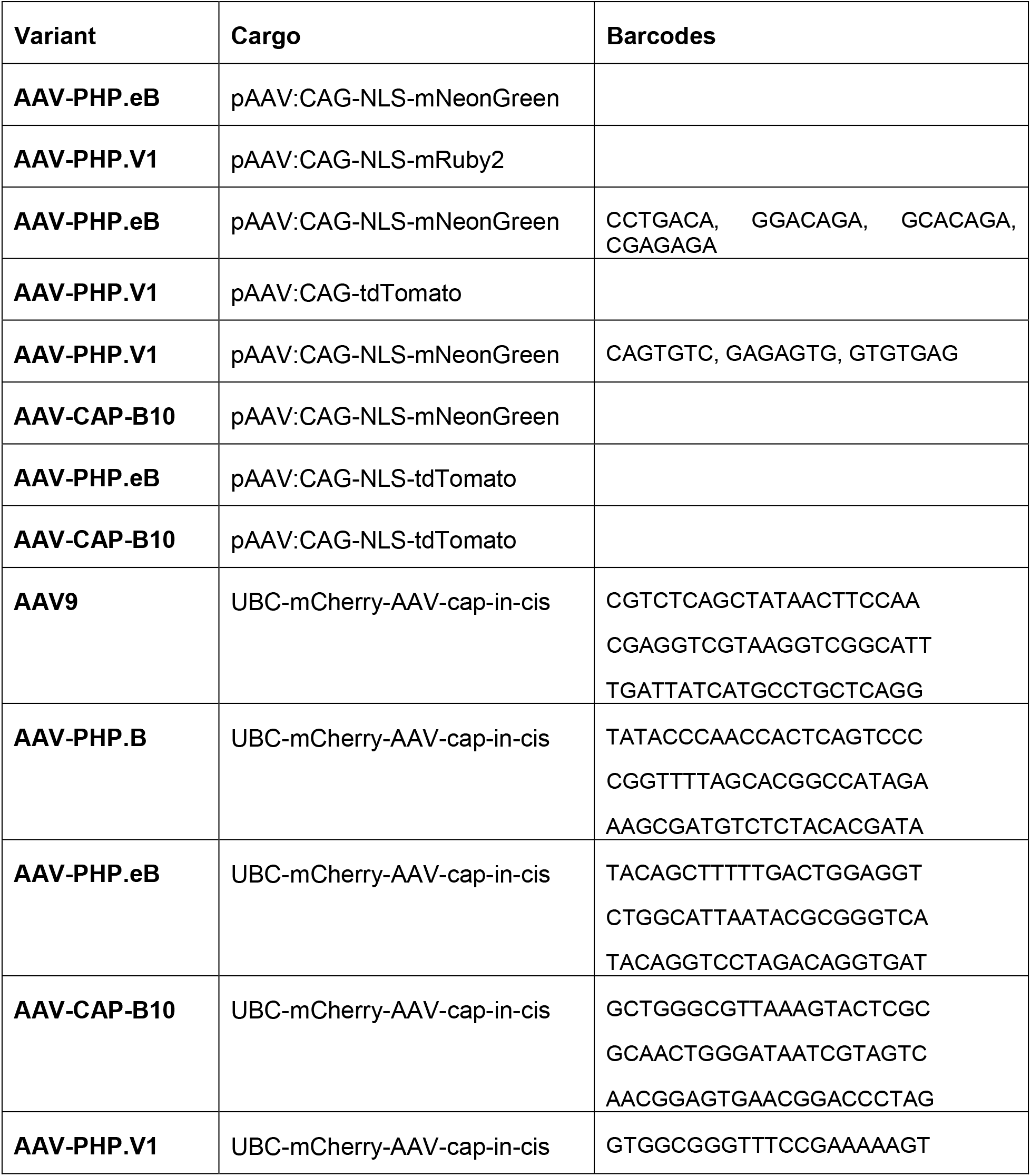
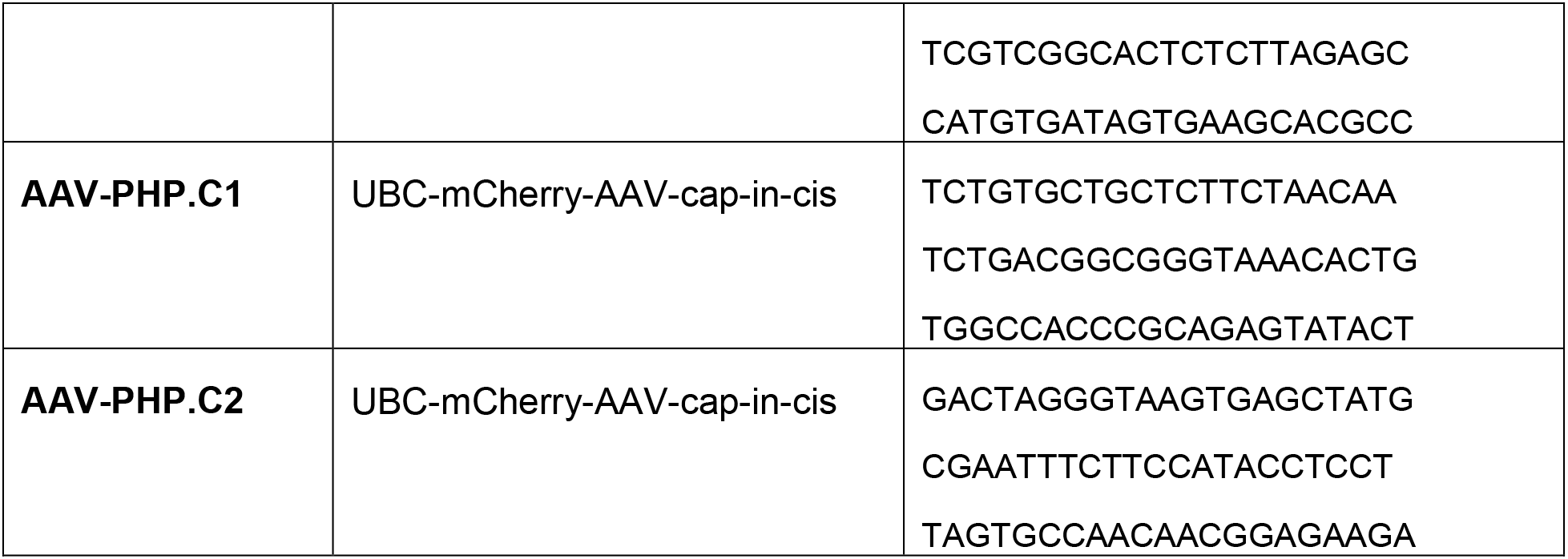
Variant Barcodes.

**Supplemental Table 6.** Differentially Expressed Genes. Supplemental file contains one tab for astrocytes, pericytes, and OPCs, with the following fields.

| Field Name | Description |
| --- | --- |
| **Gene ID** | The Ensembl Gene ID |
| **Gene name** | The canonical gene name |
| **P Non Zero** | The probability of a cell expressing this gene in this cluster |
| **P Non Zero Rest** | The probability of a cell expressing this gene in the other cell subtype clusters |

**Supplemental Table 7.** Differentially Expressed Genes Across Time Points. Supplemental file contains one tab per cell type, with the following fields.

| Field Name | Description |
| --- | --- |
| **Gene ID** | The Ensembl Gene ID |
| **Gene name** | The canonical gene name |
| **Mean expression** | The mean expression of this gene in this group |
| **L2FC** | The log fold change of this gene |
| **L2FC SE** | The standard error of the L2FC |
| **Stat** | The stat, as reported by DESeq2 |
| **P-value** | The unadjusted P-value |
| **Adjusted P-value** | The adjusted P-value |

